# BART-spatial unravels biologically significant transcriptional regulators from spatial omics data

**DOI:** 10.64898/2026.05.05.723027

**Authors:** Jingyi Wang, Hongpan Zhang, Zhenjia Wang, Chongzhi Zang

## Abstract

Transcriptional regulators (TRs) are crucial regulators of cell fate decisions by activating or repressing lineage-specific genes and integrating environmental signals with intrinsic networks. Identifying functional TRs is essential for understanding development, tissue organization, and disease. Emerging spatial transcriptomics and epigenomics technologies now provide near-single-cell resolution mapping of genomic features while preserving information of each cell’s physical location and microenvironment which influence TR activity. Despite these advances, identifying active TRs in spatial data remains challenging due to low TR expression and the fact that TR activity often does not correlate directly with mRNA levels. Moreover, existing tools mainly designed for non-spatial single-cell data overlook spatial heterogeneity. To bridge this gap, we developed BART-spatial (Binding Analysis for Regulation of Transcription for spatial omics data), an innovative computational method to infer functional TRs from spatial omics data. BART-spatial integrates spatial variability and pseudotemporal information with publicly available TR binding profiles. Applied to multiple spatial datasets from diverse platforms, including 10x Visium, Visium HD, Atera, and spatial RNA-ATAC-seq, BART-spatial consistently outperforms existing methods, identifying state-specific TRs and revealing regulators undetectable by expression alone. Its compatibility with spatial epigenomics data further strengthens its utility and enables cross-validation. Overall, BART-spatial provides a powerful and robust tool for decoding spatially resolved gene regulatory programs.

## Introduction

Transcriptional regulators (TRs), including transcription factors (TFs) and chromatin regulators, play key roles in regulating gene expression and determining cell identity during cell differentiation^1–3^. By interacting with genomic DNA and chromatin, TRs can activate or repress lineage-specific genes and integrate extrinsic environmental signals with intrinsic transcriptional networks to guide cell fate decisions^4^. Therefore, accurately identifying functional TRs is crucial for understanding the regulatory mechanisms in many biological processes, including development, tissue formation, and disease progression.

Over the past decades, high-throughput genomics technologies have empowered cellular and molecular studies of complex and heterogeneous biological systems. For example, single-cell sequencing-based genomic assays, such as scRNA-seq and scATAC-seq, have enabled profiling of gene expression and chromatin accessibility profiles, respectively, at the resolution of individual cells. These techniques allow researchers to dissect cellular heterogeneity and investigate gene regulation in different biological processes. Although each cell is captured at a static snapshot in time, computational methods, such as Monocle3^5^, can order cells along pseudotime based on their genomic-profile similarities, and generate a trajectory that potentially indicates the sequence of cellular transitions.

While single-cell-based methods are powerful, they lack consideration of the spatial context in which cells exist. In recent years, spatial transcriptomics technologies have emerged, enabling gene expression profiling with spatial location information^6^. Using either sequencing or imaging, high-resolution spatial transcriptomics techniques such as 10x Genomics Visium HD, Xenium, and Atera can achieve single-cell or subcellular resolution, making it possible to map gene expression at fine scales. In parallel, spatial epigenomics methods such as spatial ATAC-seq^7,8,9^ can also achieve single-cell resolution and offer a direct readout of chromatin profiles that potentially indicate regulatory activities. These spatial omics platforms preserve the physical location of each cell or spot within a tissue section, offering an opportunity to investigate how tissue architecture and local microenvironment may influence gene regulation and shape cellular identity and function.

Identification of functional TRs from spatial omics data remains particularly challenging. Although spatial transcriptomics is now widely used and spatial epigenomics techniques are emerging, no current approach can directly measure TR binding profiles with spatial resolution. Spatial ATAC-seq can provide chromatin accessibility information, but its ability to infer TR activity remains indirect. There is a critical need for computational methods that can infer functional TR activity from spatial transcriptomics data alone. Moreover, a TR’s expression level measured by spatial transcriptomics data may be insufficient to infer its regulatory activity, because TR activity is often modulated post-transcriptionally and does not always correlate with its mRNA level^10^, especially in spatial contexts where signaling gradients and microenvironments can influence TR activation^4,11^. In addition, many TRs are expressed at low levels, making it difficult to distinguish active regulators based on expression alone^12^. Most existing computational methods for TR prediction are primarily designed for bulk or single-cell data and do not explicitly account for spatial organization. Integrating spatial variability with biological progression adds another layer of complexity, as few methods are equipped to capture region-specific or spatially dynamic regulatory activity.

Several computational tools have been developed for TR prediction from spatial transcriptomics data but limitations remain. For example, SpaTrack^13^ constructs spatially informed trajectories and infers pseudotime-dependent features. It then uses co-expression between predefined TRs and genes to generate a ranked list of TR–gene–weight triplets as output. In contrast, SCRIPro^14^ is specifically designed to infer TR activity from single-cell and spatial omics data at the cell group level (“SuperCell”), without inferring a trajectory or generating spatially variable features. It then predicts TRs by incorporating both TR expression and functional impact via an *in silico* deletion approach^15^.

To address these challenges and gaps in current methodology, we introduce BART-spatial (Binding Analysis for Regulation of Transcription for spatial omics data), a computational method to infer functional TRs from spatially resolved transcriptomic or epigenomic data. BART-spatial integrates both spatial variability and pseudotemporal dynamics of gene expression or open chromatin information to generate biologically informed predictions of TR activity. It leverages genome-wide TR binding profiles from publicly available ChIP-seq data to enhance prediction accuracy, even in the absence of high TR expression. We demonstrate that BART-spatial accurately identifies state-specific TRs across 5 spatial omics datasets, covering diverse platforms including Visium, Visium HD, Atera, spatial RNA-seq, and spatial ATAC-seq, and different biological systems including human cancer, mouse small intestine, and mouse embryo. We show that BART-spatial outperforms existing tools in recovering known functional, state-specific TRs. Furthermore, we demonstrate BART-spatial’s compatibility with spatial epigenomics data, enabling robust prediction of TR activity from chromatin accessibility landscapes and facilitating cross-modal validation between transcriptomic and epigenomic data. Together, our work establishes BART-spatial as a generalizable framework for linking spatial organization, biological trajectories, and transcriptional regulation.

## Results

### BART-spatial integrates spatial and pseudotemporal features with public ChIP-seq data to predict functional TRs

BART-spatial is designed for predicting functional transcriptional regulators (TRs) from spatial omics data (Figure 1). TR prediction is performed using the BART algorithm^16^, which leverages a large collection of public ChIP-seq data that measure TRs’ genome-wide binding profiles. BART-spatial assumes that the input features share similar regulatory patterns (co-regulated or bound by same TRs). In the context of spatial omics data, the input features are expected to exhibit variability in spatial distribution, commonly known as spatially variable features (SVFs, e.g., spatially variable genes). In BART-spatial, SVFs are identified based on Moran’s I^17^, a spatial autocorrelation statistic that quantifies the extent to which similar values cluster in space. A positive deviation of Moran’s I from expectation indicates non-random spatial organization, suggesting high localized regulatory activity.

**Figure 1.**
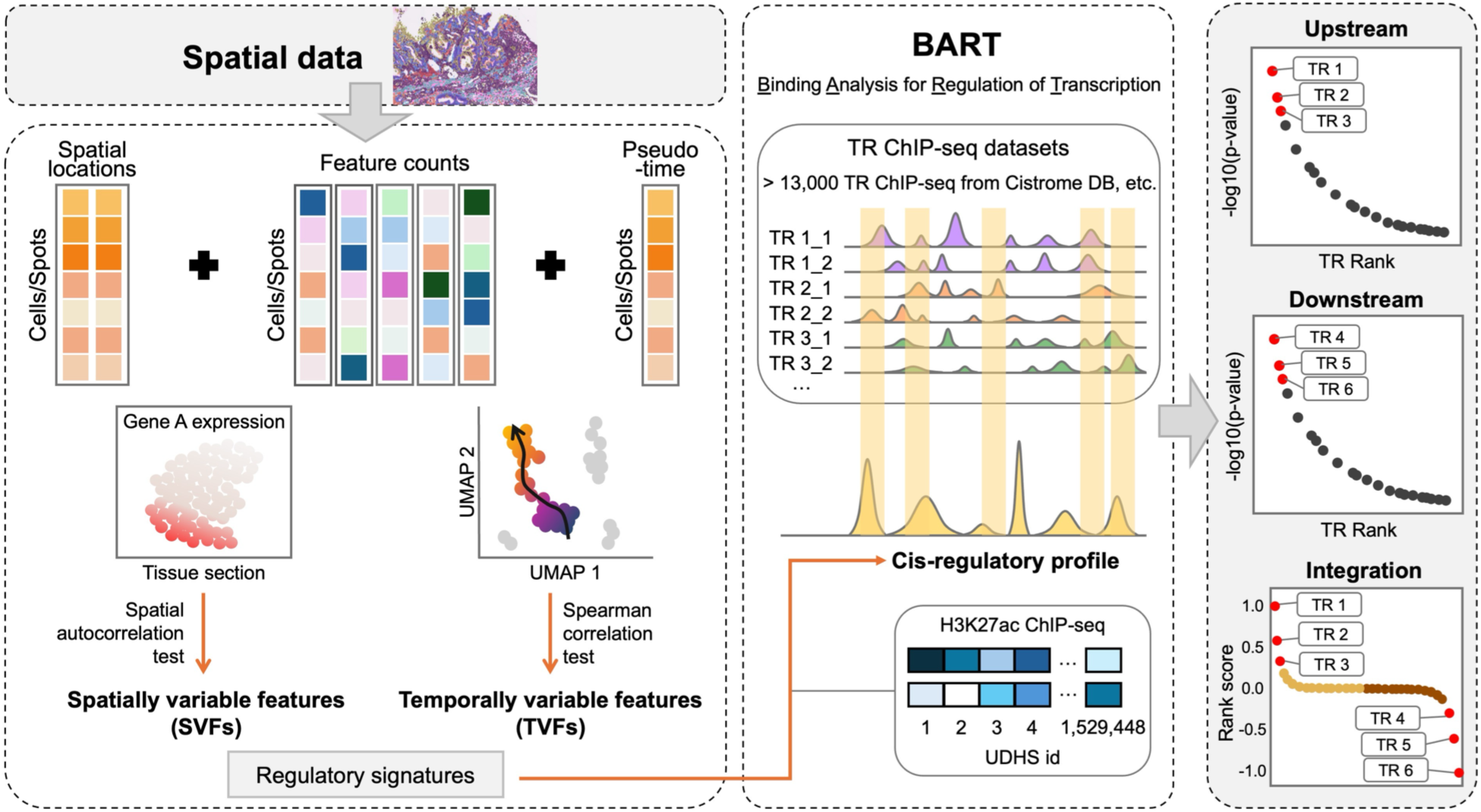
Schematic of BART-spatial. BART-spatial takes a spatial omics dataset (e.g. spatial transcriptomics) as input. It extracts spatially variable features (SVFs) based on spatial autocorrelation test by Moran’s I. In parallel, it constructs a trajectory rooted in a user-specified starting cell state, modeling cellular transitions over pseudotime, and uses Spearman correlation test to identify temporally variable features (TVFs). SVFs and TVFs are then integrated and used as input for the BART algorithm to generate a cis-regulatory profile, which is then aligned with TR ChIP-seq datasets. Sttistical significance is assessed using a Wilcoxon rank-sum test, and TRs are ranked by integrating AUC, p-value, and Z-score. The final output is ranked lists of upstream-and downstream-active TRs along the trajectory of interest.

Because spatial variability is usually evaluated for each feature independently, SVFs only reflect each feature’s difference in spatial distribution but do not necessarily represent a shared regulatory program among features. Thus, SVFs alone may not fully satisfy BART’s assumption for coordinated regulation among input features. To address this limitation, we introduced temporally variable features (TVFs), which capture coordinated changes associated with a common biological context or dynamic process. Specifically, BART-spatial constructs a trajectory rooted in a user-specified starting point, modeling cellular transitions over pseudotime. Alternatively, users can provide a precomputed cell-by-pseudotime matrix as the pseudotemporal trajectory input. BART-spatial then uses Spearman correlation to identify features (gene expression or chromatin accessibility) with consistent changes along the trajectory and defines them as TVFs. By intersecting TVFs and SVFs, BART-spatial prioritizes features that are both spatially structured and pseudotemporally dynamic, enhancing the biological specificity of the input feature set (Figure S1). This refined feature set is then used as the regulatory signature input for the BART algorithm to predict TRs that function in the biological events.

For the RNA mode, the BART algorithm first infers a cis-regulatory profile, represented as a ranked list of union DNase I hypersensitive sites (UDHS)^18^, from the input gene set based on public H3K27ac ChIP-seq data through an adaptive Lasso-based semi-supervised learning approach^19^. It then compares this profile to over 13,000 known TR ChIP-seq datasets. Each TR profile is projected onto the UDHS regions as a binary vector, and its similarity to the cis-regulatory profile is quantified using area under the ROC curve (AUC) analysis. TR predictions are based on a series of statistical tests and rank integration.

For the ATAC mode for spatial epigenomics data, BART-spatial requires a chromatin accessibility matrix as input instead of a gene expression matrix for spatial transcriptomics data. In the ATAC mode, BART-spatial uses the union of SVFs and TVFs, ranking the combined set from most to least significant. To ensure accurate state-specific inference, we require more than 50 genomic regions overlapping between state-specific TVFs and SVFs. Different from the RNA mode, BART-spatial-ATAC bypasses the adaptive Lasso-based semi-supervised learning step and directly maps input regions to UDHS. A regulatory profile is generated by scoring UDHS regions based on their overlap with accessible chromatin regions, and the resulting profile undergoes the same steps as in the RNA mode for TR prediction.

### BART-spatial identifies state-specific TRs along HPV-negative oral squamous cell carcinoma progression from Visium data

To evaluate the performance, we first applied BART-spatial to a spatial transcriptomics dataset generated using 10x Genomics Visium, one of the most widely adopted spatial transcriptomics platforms^20^. This publicly available dataset generated by Aurora and Cao *et al.*^20^ is from a sample of HPV-negative oral squamous cell carcinoma (OSCC), the most prevalent subtype of head and neck squamous cell carcinoma (HNSCC) and a well-characterized model for transcriptional regulation ^20,21^. This dataset profiled spatially resolved transcriptomes spanning three tumor regions: tumor core (TC), transitory region, and leading edge (LE) (Figure 2a). We used the spatial trajectory of tumor differentiation from TC to LE, as described in the original study^20^, as a benchmark to assess BART-spatial’s ability to capture region-specific and progression-related transcriptional regulatory activities, compared with another two methods, SCRIPro and SpaTrack. As a result, both BART-spatial and SpaTrack successfully inferred temporal trajectories consistent with the observed spatial progression of OSCC from TC to LE (Figure 2b, c). The spatial patterns of expression levels of several established region-specific marker genes, such as *LCN2* for TC^20,22^ (Figure 2d), *KRT6C* for the transitory zone^20,23^ (Figure 2e), and *COL1A2* for LE^20^ (Figure 2f), further supported the validity of the inferred trajectories.

**Figure 2.**
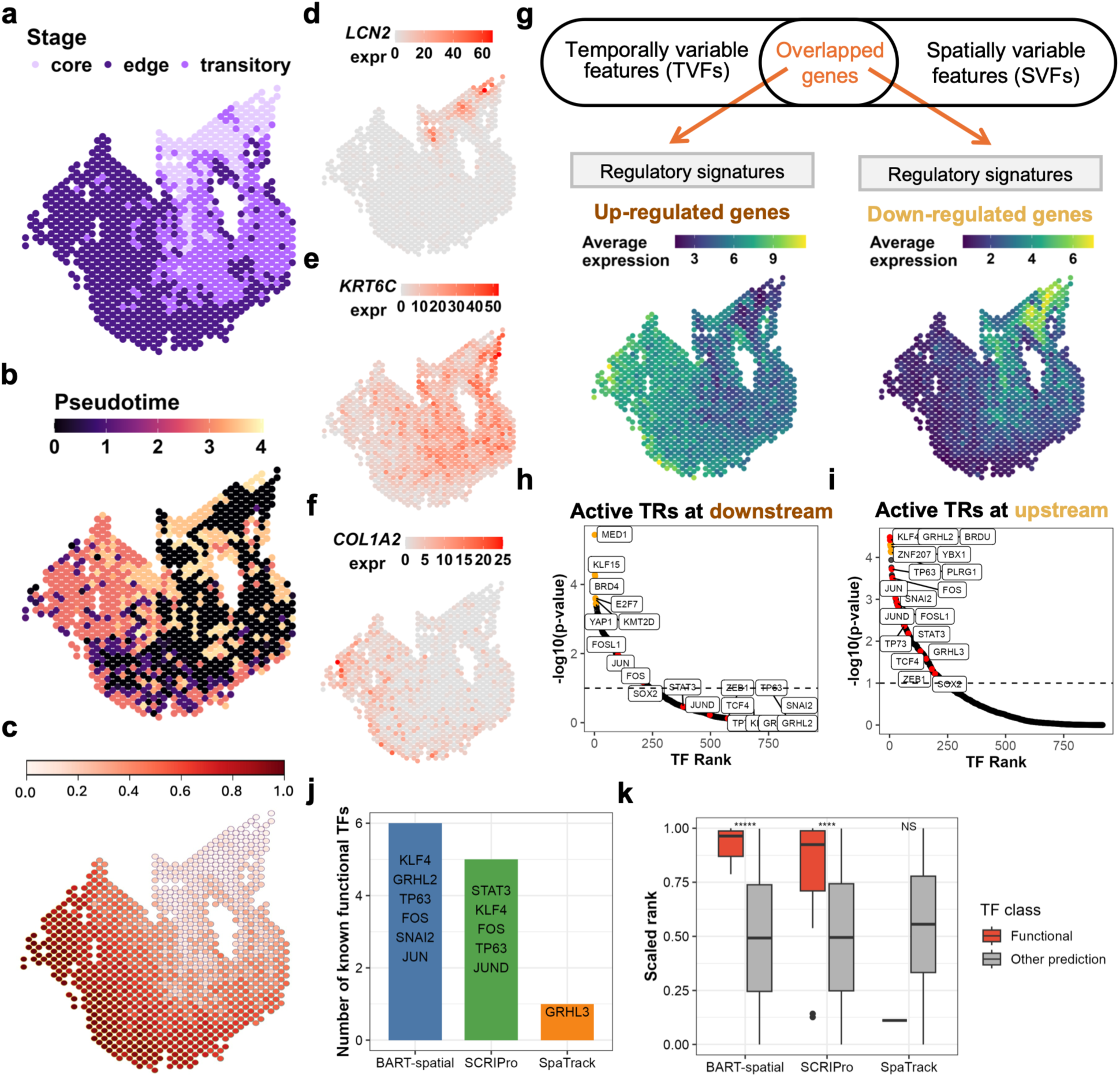
BART-spatial application on 10x Visium data for HPV-negative oral squamous cell carcinoma. (a) Spatial distribution of tumor core (TC), leading edge (LE), and transitory regions in HPV-negative OSCC sample. (b) Pseudotime analysis by BART-spatial from TC to LE visualized on the spatial map. (c) Pseudotime analysis by SpaTrack from TC to LE visualized on the spatial map. (**d**-**f**) Spatial map visualization of the expression of the key genes *LCN2* (**d**), *KRT6C* (**e**), and *COL1A2* (**f**). (**g**) BART-input gene sets are separated into upregulated along pseudotime (left, as input genes for predicting downstream-active TRs) and downregulated along pseudotime (right, as input genes for upstream-active TRs). The averaged expression of input genes is shown on the spatial map. (**h**, **i**) BART-spatial prediction results for downstream-active TRs (**h**) and upstream-active TRs (**i**). Highlighted in yellow are the top 6 TRs predicted by BART-spatial and highlighted in red are known marker TRs in OSCC development with literature support. (j) The numbers of known functional TRs in OSCC development among the top 25 TRs predicted by BART-spatial, SCRIPro, and SpaTrack. (k) The scaled ranks of known functional TRs against all the other predicted TRs in the upstream prediction results by BART-spatial, SCRIPro, and SpaTrack. *****, p < 10^-5^; ****, p < 10^-4^; NS, non-significant, by one-sided Wilcoxon test.

We then sought to detect regulatory signatures (differentially expressed genes along the trajectory) from this dataset. Along the differentiation trajectory from TC to LE, we identified 48 upregulated and 302 downregulated genes with BART-spatial (Figure 2g). We focused on upstream-active TRs, which are more active in TC, inferred from downregulated genes along the differentiation trajectory. To assess biological relevance, we curated a benchmark list of 15 TRs, each of which is known to function in OSCC or HNSCC based on literature with experimental support (GRHL3^20^, TCF4^20^, TP63^24,25^, TP53^26^, GRHL2^27^, SOX2^28^, KLF4^29^, TP7^30^, SNAI2^31^, ZEB1^32^, FOSL1^33^, STAT3^34^, FOS^33^, JUN^33^, and JUND^33^). We observed that all of these 15 TRs were highly ranked in the BART-spatial upstream prediction results and reached statistical significance (p-value < 0.1) (Figure 2h, i). Among the top 25 upstream-active TRs predicted by each of the three methods, BART-spatial successfully recovered 6 of the 15 known functional TRs, compared to 5 identified by SCRIPro and 1 by SpaTrack (Figure 2j), indicating the superior performance of BART-spatial than the other two methods.

We next evaluated whether the known functional TRs were scored and ranked higher than all the other TRs in each method’s prediction. To achieve this, we rescaled the output TR ranks from each method to a 0–1 range, with 1 representing the highest predicted activity. In both BART-spatial and SCRIPro results, the known functional TRs exhibited significantly higher ranks than the other TRs (*P* = 1.25×10^-8^ for BART-spatial, and 9.05×10^-5^ for SCRIPro, respectively, Figure 2k). This distinction was not observed in the SpaTrack output. Notably, all 15 known TRs had scaled ranks above 0.75 in the BART-spatial predictions, whereas their ranks were more dispersed in the SCRIPro results with two lowly-ranked outliers (Figure 2k).

In summary, we demonstrated that BART-spatial outperformed other methods in inferring upstream-active TRs along the tumor progression trajectory. We were able to identify more biologically relevant TRs with consistently high scores or ranks by BART-spatial. These results highlight BART-spatial’s sensitivity and functional relevance for TR prediction from spatial transcriptomics data.

### BART-spatial identifies state-specific TRs in enterocyte maturation from Visium HD data

To assess the versatility of BART-spatial across different platforms, biological systems, and species, we next applied BART-spatial to a Visium HD spatial transcriptomics dataset for mouse small intestine from the 10x Genomics website. The mouse intestine is a well-established model for studying cell development due to its well-defined spatial architecture along the crypt–villus axis. Intestinal stem cells reside at the base of the crypts, where they continuously proliferate and give rise to transient amplifying progenitors^35^. As these progenitor cells migrate upward toward the villus, they differentiate into various mature cell types^35^. We obtained cell type annotations from Haber et al.^36^, and labeled enterocytes, Paneth cells, goblet cells, stem cells, and endocrine cells on the spatial map (Figure 3a). Among them, enterocytes, which are hyperpolarized epithelial cells responsible for nutrient absorption, are particularly abundant^35^. Focusing on the enterocyte lineage, we observed a clear spatial progression from progenitor cells in the crypt-villus junction to mature enterocytes along the villus axis (Figure 3b). This developmental trajectory was further supported by pseudotime analysis, which captured the temporal ordering of cells from immature to mature states (Figure 3c). The spatial expression patterns of several enterocyte marker genes, including *Krt19*^37,38^ (Figure 3d), *Anpep*^39^ (Figure 3e), *Maf*^40^ (Figure 3f), and *Clca4a*^41^ (Figure 3g), aligned with this progression, further validating the inferred trajectory.

**Figure 3.**
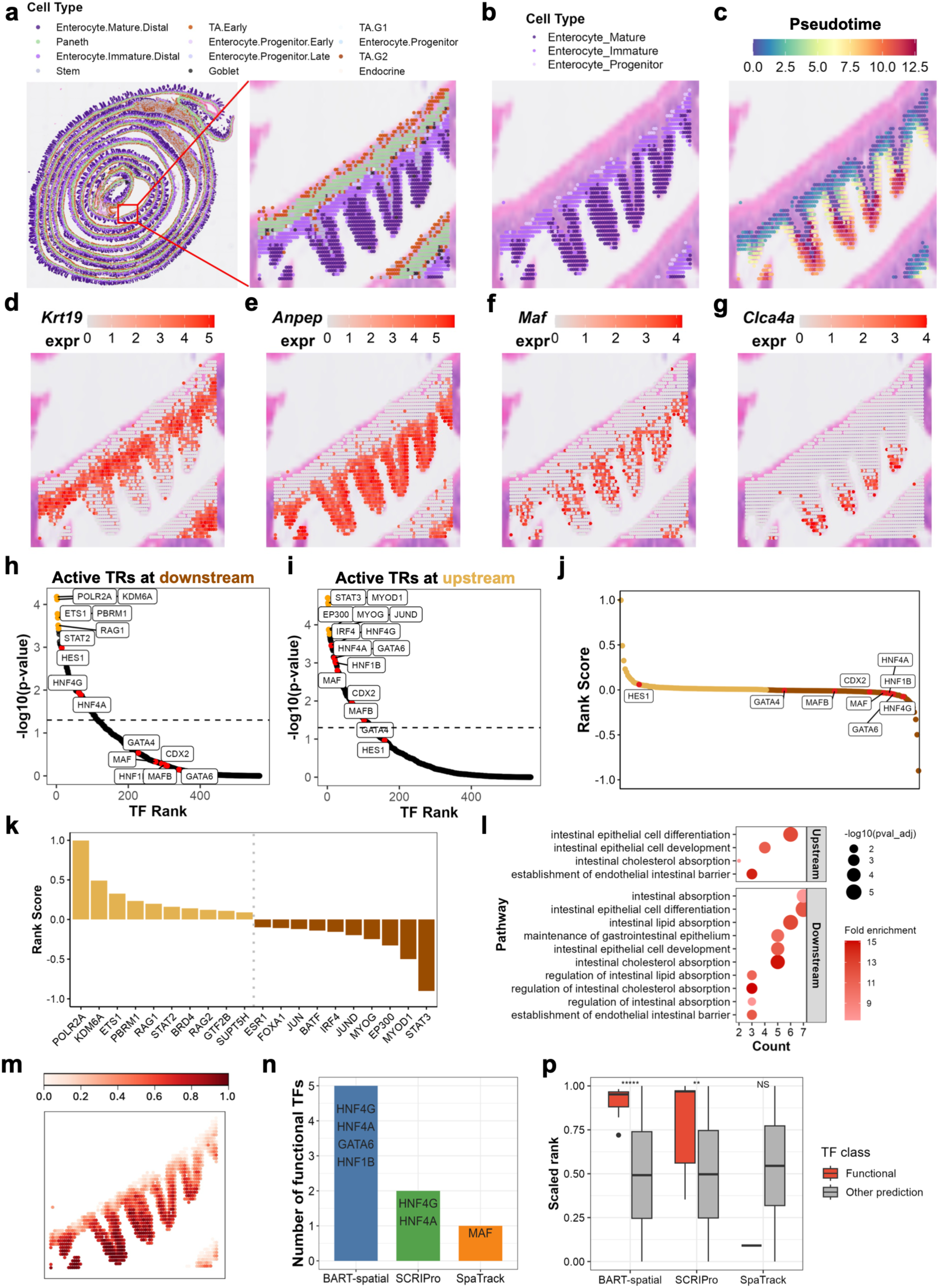
BART-spatial identifies state-specific TRs in enterocyte maturation from Visium HD data. (a) Spatial distribution of all cell types in the mouse small intestine. (b) Spatial distribution of enterocyte stages in focused regions of the mouse small intestine. (c) Pseudotime analysis by BART-spatial from enterocyte progenitor to mature enterocyte visualized on the spatial map. (**d**-**g**) Spatial map visualization of the expression of the key genes *Krt19* (**d**), *Anpep* (**e**), *Maf* (**f**), and *Clca4a* (**g**). (**h**, **i**) BART-spatial results for downstream-active TRs predicted from upregulated genes (**h**) and upstream-active TRs predicted from downregulated genes (**i**). Highlighted in yellow are the top 6 TRs in BART-spatial prediction results. TRs highlighted in red are known marker TRs in the enterocyte maturation process with supporting literature. (j) Integration of downstream and upstream predictions, highlighting the known functional TRs. (k) Integration of downstream and upstream predictions, showing the top 10 TRs from each of the downstream and upstream TR lists. (l) GO enrichment analysis for the putative target genes of BART-spatial-predicted TRs. (m) Pseudotime analysis by SpaTrack from enterocyte progenitor to mature enterocyte visualized on the spatial map. (n) The numbers of known functional TRs in enterocyte maturation among the top 25 TRs predicted by BART-spatial, SCRIPro, and SpaTrack. (**p**) The scaled ranks of known functional TRs against all the other predicted TRs in the downstream prediction results by BART-spatial, SCRIPro, and SpaTrack. *****, p < 10^-5^; ****, p < 10^-4^; NS, non-significant, by one-sided Wilcoxon test.

By integrating both spatial and temporal information, BART-spatial identified 1,923 significant genes, including the known marker genes mentioned above, as feature input for predicting functional TRs involved in enterocyte development. The identified genes were further stratified based on their expression dynamics: upregulated genes along pseudotime were treated as downstream-active genes, which were used to infer TRs active during late development; downregulated genes along pseudotime were treated as upstream-active genes, which were used to predict TRs active in early developmental states. Using this approach, BART-spatial successfully identified key TRs known to regulate enterocyte development, including HNF4G^42^, HNF4A^43^, HNF1B^43^, GATA6^44^, GATA4^44^, MAF^40^, MAFB^40^, CDX2^45^, and HES1^40,46^ (Figure 3h-j). Notably, the spatial activity of these TRs varied along the inferred developmental trajectory. For example, HES1 showed significant regulatory activity only in the upstream region (p < 0.05), suggesting its involvement in early progenitor states (Figure 3h). In contrast, GATA4, GATA6, MAF, MAFB, CDX2, and HNF1B were identified as downstream-active TRs, with significant activity only in later stages of enterocyte maturation (p < 0.05) (Figure 3i). Among these, MAF is known as an enterocyte-specific TF critical for differentiation along the villus axis and *Maf* is expressed in mature enterocytes but largely absent in progenitor cells^40^. This spatial expression pattern was also observed in the Visium HD dataset (Figure 3f), supporting BART-spatial’s prediction. Additionally, some of the top 10 downstream-active TRs, including STAT3^47,48^, JUND^49,50^, EP300^51^, IRF4^52^, and BATF^53,54^, have been reported in processes related to wound healing and immune barrier formation (Figure 3k), suggesting that BART-spatial not only captures trajectory-related regulators but also identifies context-specific TRs involved in broader biological processes.

To further evaluate the functional relevance of the predicted TRs, we performed Gene Ontology (GO) enrichment analysis on their putative target genes identified using the TRRUST database^55^. The target genes of downstream-active TRs were highly enriched in biological processes associated with intestinal lipid and cholesterol absorption, which are hallmark functions of mature enterocytes. In contrast, targets of upstream-active TRs showed relatively less enrichment in nutrient uptake (Figure 3l). Taken together, these findings demonstrate that BART-spatial not only finds TRs involved in enterocyte development but also correctly infers their temporally specific regulatory activity.

Similarly, we benchmarked BART-spatial against SpaTrack and SCRIPro using this dataset. SpaTrack inferred a trajectory that was consistent with the BART-spatial-derived enterocyte maturation trajectory (Figure 3m). Among the top 25 upstream-and downstream-active TRs predicted by each method, we assessed whether they include nine literature-supported TRs, which are HNF4G, HNF4A, HNF1B, GATA6, GATA4, MAF, MAFB, CDX2, and HES1. BART-spatial identified four of these TRs, SCRIPro recovered two, and SpaTrack only found MAF in its downstream predictions (Figure 3n). Next, we evaluated the scaled ranks of these known TRs relative to the other TRs in the prediction results from each method. In the BART-spatial results, eight of the nine TRs exhibited scaled ranks above 0.75 (top 25%) and were significantly higher than those of other TRs. While SCRIPro also showed a significant difference in the scaled ranks between known TRs and the rest, the distribution of the known TRs was more scattered. In contrast, MAF did not rank higher than other predicted TRs in the SpaTrack output (Figure 3p). These benchmarking analyses further support the superior performance of BART-spatial.

### BART-spatial reveals TRs functional in two states of cancer-associated fibroblast from Atera data

We next applied BART-spatial to a spatial transcriptomics dataset for human breast cancer tissue generated by the recently released 10x Genomics Atera, a single-cell-resolution, whole-transcriptome spatial profiling platform. Cell-type annotation of this dataset identified two subtypes of cancer-associated fibroblasts (CAFs): ductal carcinoma in situ (DCIS)-associated and invasive-associated. These two types of CAF populations occupied partially overlapping spatial regions (Figure 4a, b), implying a continuous cell-state transition rather than completely independent populations. DCIS represents a non-invasive breast cancer stage in which abnormal epithelial cells remain confined within the ducts. Based on this biological context and the spatial distribution of the two CAFs, we hypothesized a trajectory from DCIS-associated CAFs to invasive-associated CAFs (Figure 4c).

**Figure 4.**
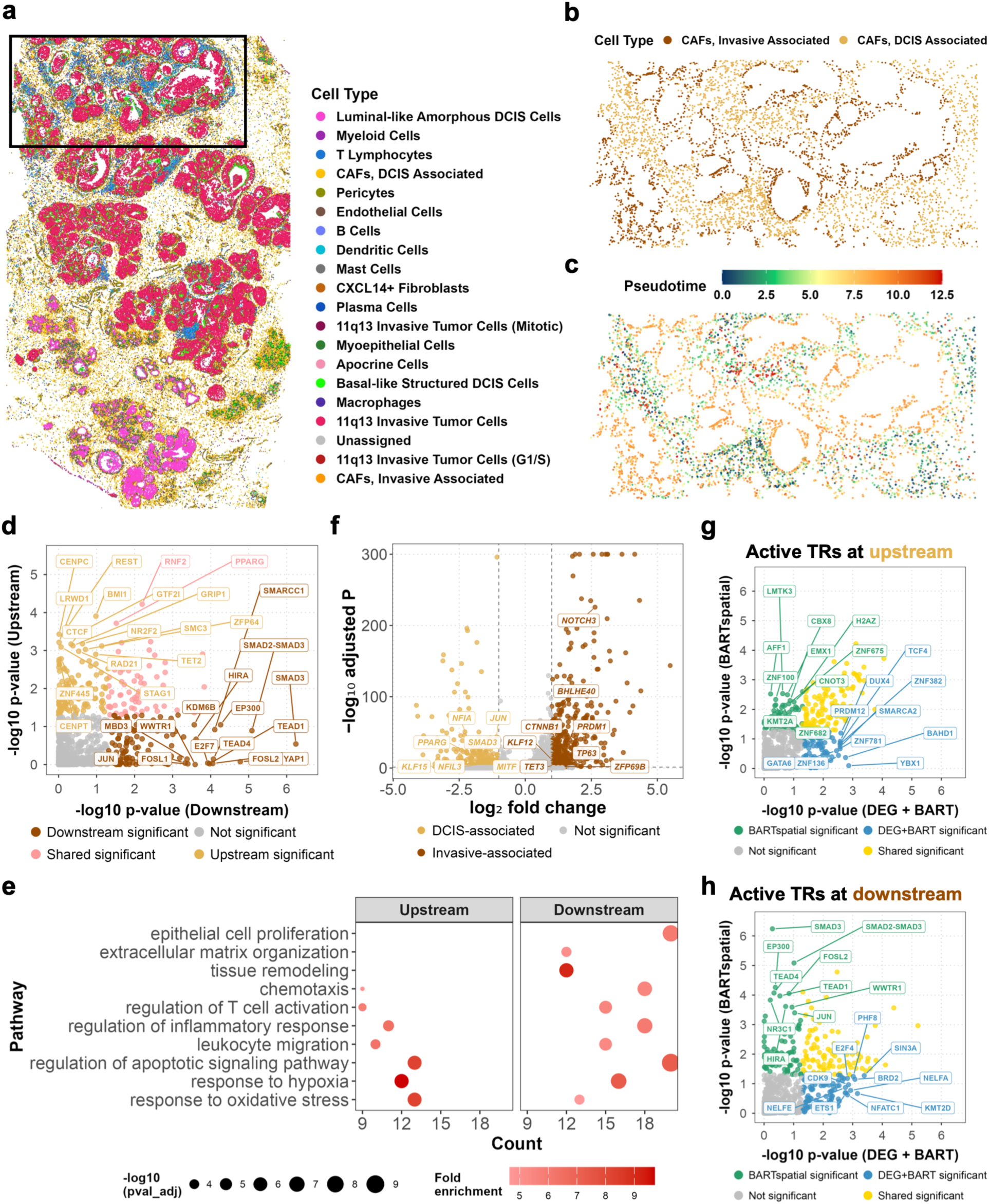
BART-spatial reveals invasive CAF progression from single-cell-resolution, whole-transcriptome Atera data. (a) Spatial distribution of all cell types in the human breast cancer. (b) Spatial distribution of CAF states in focused regions. (c) Pseudotime analysis by BART-spatial from DCIS-associated CAFs to invasive-associated CAFs visualized on the spatial map. (d) Scatter plot showing BART-spatial downstream and upstream predictions. Top 15 predicted TRs in each side were labeled. (e) GO enrichment analysis for the putative target genes of BART-spatial-predicted TRs. (f) Volcano plot showing differential gene expression between invasive-associated and DCIS-associated CAFs. Genes encoding BART-spatial-predicted significant TRs are labeled. (**g**, **h**) Comparison of upstream-active (**g**) and downstream-active (**h**) TRs identified by BART-spatial and canonical DEG+BART analysis. TRs significant only by BART-spatial, only by DEG+BART, by both methods, or by neither method are indicated by different colors.

Using this trajectory, we identified two sets of TRs active in the upstream DCIS-associated state and the downstream invasive-associated state, respectively (Figure 4d, S2a). Upstream regulators included factors such as RNF2^56,57^, PPARG^58^, REST^59^, and CTCF^60^, suggesting broader regulatory programs associated with cell state maintenance and chromatin-level gene regulation in breast cancer. In contrast, downstream regulators included SMAD-family members^61,62^, YAP/TEAD^63–65^, AP-1-related factors^66^, indicating a shift toward stromal remodeling and invasive tumor-associated programs, a pattern confirmed by GO enrichment analysis using the putative target genes of upstream-and downstream-active TRs (Figure 4e). These results support a coherent progression from DCIS-associated to invasive-associated CAF states, marked by a transition from general regulatory programs to invasion-associated mechanisms involved in matrix remodeling, stromal activation, and tumor progression.

Interestingly, although predicted TRs, such as PPARG, were also significantly differentially expressed between DCIS-associated and invasive-associated CAFs, changes in predicted regulatory activity for many TRs were not correlated with their expression levels (Figure 4f). This finding highlights BART-spatial’s ability to identify TRs whose spatially variable regulatory activities are not apparent from expression alone. A notable example was the YAP1/TEAD1 transcriptional complex, which was predicted to be preferentially active in invasive-associated CAFs despite neither *YAP1* nor *TEAD1* being identified as a differentially expressed gene. This prediction is consistent with the established role of YAP1/TEAD1 signaling in aggressive breast cancer and its potential as a therapeutic target in aggressive breast cancer^65^. TEAD1 has been reported to form distinct types of biomolecular condensates at inactive storage sites near heterochromatin and at transcriptionally active loci, without corresponding changes in its transcription or protein level^67^. Notably, several downstream-active regulators, including SMAD3 and stem cell-associated factors NANOG and POU5F1, were identified only by BART-spatial and not by canonical BART using differentially expressed genes as input (Figure 4g, h, S2b-f), underscoring the importance of incorporating spatially structured feature variation into TR inference.

### BART-spatial reveals TR regulatory activity not apparent from expression

We used BART-spatial to analyze a spatial omics map of E13 mouse embryos^8^, with a focus on neural development (Figure 5a). In the developing central nervous system (CNS), radial glial cells play dual roles as both progenitors of neurogenesis and guides of newly formed neuron migration^68^. Consistent with the original study, we used the spatial transcriptomics profile to identify a developmental trajectory from radial glia to postmitotic premature neurons in the spatial landscape (Figure 5b). We validated this trajectory by pseudotime inference (Figure 5c) and the expression patterns of marker genes associated with neural differentiation, including *Pax6*^8,68^ (Figure 5d), *Fabp7*^8^(Figure 5e), and *Myt1l*^8^ (Figure 5f). We then applied BART-spatial on this trajectory and identified key regulators involved in radial glia differentiation, such as PAX6^8,68^ (Figure 5g, h). Notably, BART-spatial could distinguish TRs with state-specific regulatory activity along the trajectory. For example, NEUROD2, a TF known to be active in postmitotic neurons and involved in terminal positioning during the final stages of radial glia migration^68,69^, was ranked as the top downstream-active regulator in the BART-spatial results (Figure 5g), while it was not detected as significant in the upstream TR list (Figure 5h). In addition to using a binary significance cutoff, we also obtained insights into TRs’ relative activity across developmental stages from their ranks in the BART-spatial results. For instance, SOX9 was identified as active in both upstream and downstream of the trajectory (Figure 5g, h), but with higher activity in the earlier stage (ranked #1 in upstream, #40 in downstream, respectively). This pattern aligns with SOX9’s established role as a radial glia progenitor marker^70–72^, as well as its remaining function in postmitotic neurons, where it activates the expression of genes critical for establishing neuronal polarity^70–72^.

**Figure 5.**
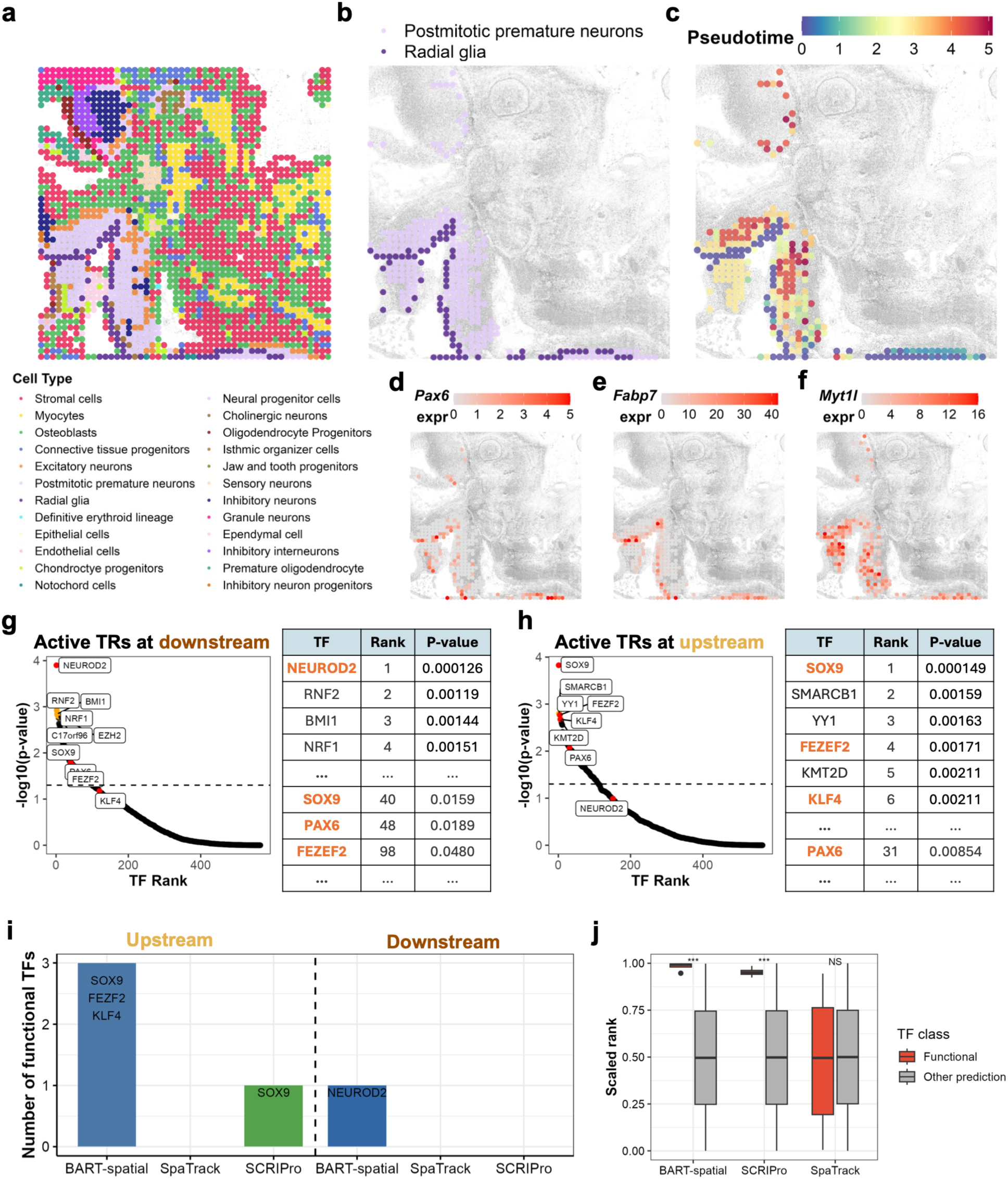
BART-spatial predicts TRs active during mouse embryonic neural development from spatial transcriptomics data. (a) Spatial distribution of all cell types in E13 mouse embryo. (b) Spatial distribution of radial glia and postmitotic premature neurons. (c) Pseudotime analysis by BART-spatial from radial glia (upstream) to postmitotic premature neurons (downstream) visualized on the spatial map. (**d**-**f**) Spatial map visualization of the expression of the key genes *Pax6* (**d**), *Fabp7* (**e**), and *Myt1l* (**f**). (**g**, **h**) BART-spatial results for downstream-active TRs predicted from upregulated genes (**g**) and upstream-active TRs predicted from downregulated genes (**h**). The top 6 predicted TRs are marked in yellow; known marker TRs in radial glia differentiation are marked in red in the plot and highlighted in orange in the table. (i) The numbers of known TRs in radial glia differentiation among the top 25 TRs predicted by BART-spatial, SCRIPro, and SpaTrack. (j) The scaled ranks of known TRs against all the other predicted TRs in the upstream prediction results by BART-spatial, SCRIPro, and SpaTrack. *****, p < 10^-5^; ****, p < 10^-4^; NS, non-significant, by one-sided Wilcoxon test.

We compared BART-spatial with SpaTrack and SCRIPro for their ability to recover known functional TRs from this mouse embryo spatial transcriptomics dataset. Among the top 25 TRs identified by each method, BART-spatial recovered 3 of 4 known upstream-active functional TRs, as well as the only known downstream-active functional TR (Figure 5i). In contrast, although SpaTrack inferred a trajectory consistent with the BART-spatial-derived neural development trajectory (Figure S4b), it failed to recover any of the known TRs (Figure 5i). SCRIPro identified only one known upstream-active TR and no known downstream-active TRs (Figure 5i). We next examined the scaled ranks of TRs predicted by each method and compared the ranks of known upstream-active TRs with those of all other TRs. All the known upstream-active TRs achieved scaled ranks close to 1 (highly ranked) in both the BART-spatial and SCRIPro results and were clearly separated from other TRs. In contrast, this distinction was not observed in the SpaTrack results (Figure 5j).

While the predicted activities of some TRs aligned well with their gene expression patterns, such as PAX6 (Figure 5d) and SOX9 (Figure 6a), both of whose coding genes were expressed higher in the upstream state, and NEUROD2 (Figure 6b), whose coding gene was expressed higher in the downstream state, this association was not observed for all predicted TRs. For example, KLF4 ranked as the 6th upstream-active TR in the BART-spatial prediction (Figure 5h, S3a) but *Klf4* showed low and sparse expression across the spatial map (Figure 6c, d). To validate whether KLF4 is a functional regulator in relevant cell types, we leveraged the chromatin accessibility information derived from the spatial ATAC modality of this spatial multiomics dataset. The trajectory inferred from spatial ATAC-seq data showed strong concordance with that derived from spatial transcriptomics data (Figure 6e, f). We used the ATAC-seq peaks containing KLF4 motifs as putative KLF4 binding sites and assessed KLF4 activity based on the chromatin accessibility levels of these peaks across the spatial map. We observed that these KLF4 motif-containing accessible-chromatin regions exhibited differential accessibility patterns that aligned well with the inferred developmental trajectory, particularly highlighting regions enriched for radial glial cells, suggesting the differential regulatory activities of KLF4 in this developmental pathway (Figure 6g-i).

**Figure 6.**
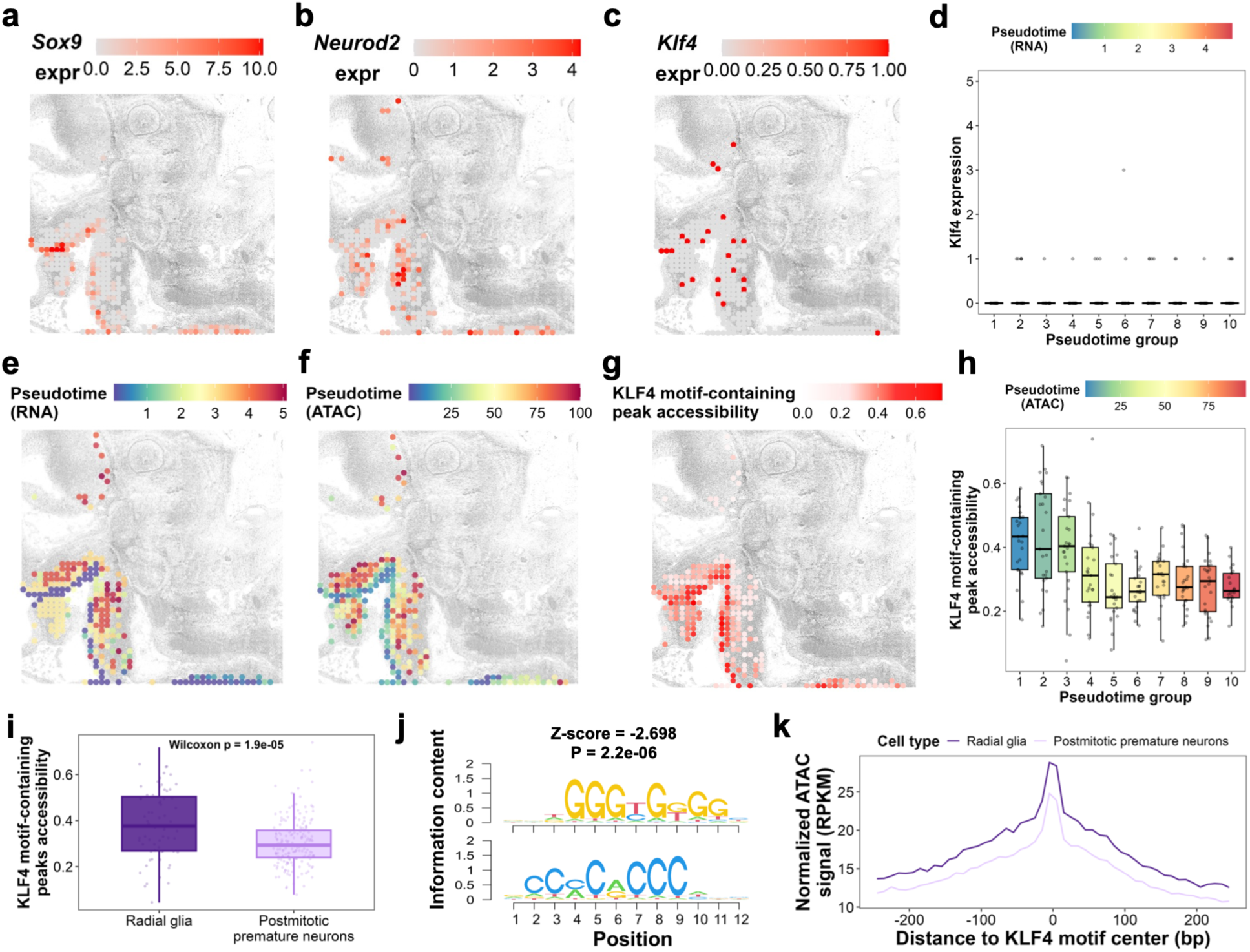
BART-spatial predicts KLF4, whose regulatory activity is not correlated with *Klf4* expression but is validated by spatial ATAC-seq. (**a**-**c**) Spatial map showing expression of *Sox9* (top 1 in upstream prediction) (**a**), *Neurod2* (top 1 in downstream prediction) (**b**), and *Klf4* (the 6^th^ in upstream prediction) (**c**). (d) *Klf4* expression across 10 sets of spots grouped by pseudotime. (**e**, **f**) Pseudotime on spatial map calculated using RNA part (**e**) and ATAC part (**f**) of the co-profiling data. (**g**, **h**) Chromatin accessibility of KLF4 motif-containing ATAC-seq peaks visualized on the spatial map (**g**) or across 10 sets of spots grouped by pseudotime (**h**), the same groups as in (**d**). (i) Chromatin accessibility of KLF4 motif-containing peaks in radial glia and postmitotic premature neurons. Each data point in a box represents the average chromatin accessibility level across all KLF4 motif-containing ATAC-seq peaks for one spot. P-value was calculated by one-sided Wilcoxon test. (j) KLF4 motif identified from differential ATAC-seq peaks with higher accessibility in radial glia. (k) Aggregate chromatin accessibility profiles centered on KLF4 motif sites within ATAC-seq peaks.

We next performed an unbiased analysis to determine whether KLF4 could be recovered directly from spatial ATAC-seq data. We generated pseudobulk ATAC-seq profiles for radial glia and postmitotic premature neurons and performed motif enrichment analysis on chromatin regions differentially accessible between the two cell states. KLF4 was independently recovered as a significantly enriched motif (Figure 6j), providing orthogonal support for its preferential association with radial glia-specific accessible chromatin. Correspondingly, aggregate pseudobulk ATAC-seq signal around KLF4 motif sites was higher in radial glia than in postmitotic premature neurons (Figure 6k).

Consistent with our observation, a previous study showed that KLF4 maintains the self-renewal capacity of neural progenitor cells and KLF4 KO promotes early neural differentiation by altering 3D chromatin architecture^29,73^. These results show that BART-spatial can identify TRs whose regulatory activity is not reflected by transcript abundance. In addition to KLF4, we validated several other BART-spatial-predicted TRs using a similar approach leveraging spatial ATAC-seq data, including CREB1^74^, LEF1^75^, MYCN^76^ and E2F3^77^ (Figure S3b-n). Therefore, BART-spatial is valuable for discovering active regulators that would be missed by co-expression-based analysis.

### BART-spatial exhibits high consistency across modality

BART-spatial is also compatible with spatial epigenomics data. Spatial ATAC-seq signals are often sparser than transcriptomics data^78^, presenting additional computational challenges. While SCRIPro and SpaTrack could not be effectively applied to spatial ATAC-seq data, BART-spatial remains applicable under its ATAC mode (Figure S4). We evaluated it on 3 spatial epigenomics datasets: the ATAC component of a spatial co-profiling dataset for E13.5 mouse embryo^8^, and two biological replicates of spatial ATAC-seq for the same system^7^ (Figure 7a-c). We focused only on the upstream-active TR predictions, as the overlap between downstream TVFs and SVFs in these data was insufficient (<50 regions).

**Figure 7.**
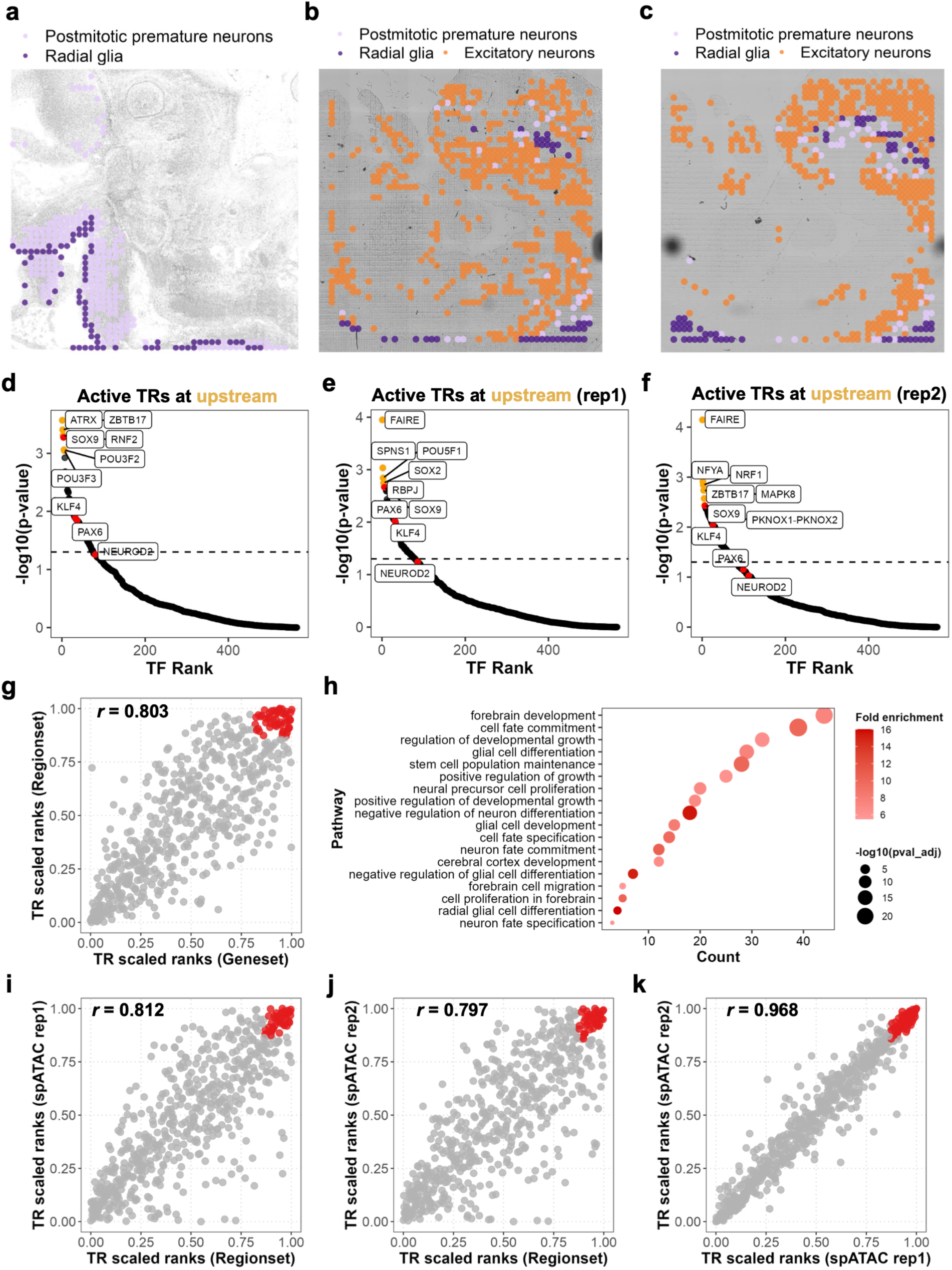
BART-spatial exhibits high consistency across modalities. (a) Spatial distribution of radial glia and postmitotic premature neurons in E13 mouse embryo spatial RNA-ATAC-seq co-profiling data. (**b**, **c**) Spatial distribution of radial glia, postmitotic premature neurons and excitatory neurons in E13 mouse embryo spatial ATAC-seq replicate 1 (**b**) and replicate 2 (**c**). (**d**-**f**) BART-spatial results for upstream-active TRs predicted from downregulated genes, using ATAC part of spatial RNA-ATAC-seq co-profiling data (**d**), spatial ATAC-seq replicate 1 (**e**), and replicate 2 (**f**). Highlighted in yellow are the top 6 TRs in BART-spatial prediction results; highlighted in red are known marker TRs in the radial glia differentiation. (g) Correlation between the scaled rank of TRs predicted using the RNA part and using the ATAC part of spatial RNA-ATAC-seq co-profiling data. Dots highlighted in red are the overlapped significant predictions (p < 0.05) between the two modalities. (h) GO enrichment analysis for the putative target genes of BART-spatial-predicted TRs. (**i**-**k**) Correlation between the scaled rank of TRs predicted using the ATAC part of spatial RNA-ATAC-seq co-profiling and spatial ATAC-seq replicate 1 (**i**), the ATAC part of spatial RNA-ATAC-seq co-profiling and spatial ATAC-seq replicate 2 (**j**), replicate 1 and replicate 2 of spatial ATAC-seq (**k**). Dots highlighted in red are the overlapped significant predictions (p < 0.05) between the two modalities.

BART-spatial’s predicted upstream-active TRs show strong concordance with previous findings, robustly recovering known functional regulators such as SOX9 and KLF4, whereas NEUROD2 was not significant in any dataset (Figure 7d-f). TR scaled ranks inferred from spatial ATAC-seq data were highly correlated with those from spatial RNA-seq data, with the correlation coefficient exceeding 0.8 (Figure 7g). GO enrichment analysis of the putative target genes of predicted TRs further supported their roles in mouse radial glia differentiation (Figure 7h). Although the spatial tissue layouts are different across datasets (Figure 7a-c), the scaled ranks of predicted TRs remained highly consistent across independent spatial ATAC-seq datasets from the same biological system (Figure 7i-k). Together, these results demonstrate that BART-spatial can generalize across spatial transcriptomics and epigenomics modalities, providing a robust, reproducible, and broadly applicable framework for identifying functional TRs from diverse spatial omics data.

## Discussion

In this study, we presented BART-spatial, a computational framework developed to infer functional TRs from spatial transcriptomics data, with an extended capability to handle spatial epigenomics data through an ATAC mode. BART-spatial integrates spatial variability with pseudotemporal dynamics to generate biologically meaningful predictions of TR activities. Across datasets, BART-spatial demonstrated superior performance, benefiting from its integration of spatial and pseudotemporal information with TR binding prior knowledge. Unlike conventional methods that rely on TR expression levels alone, BART-spatial considers the unique spatial organization of cells, making it possible to uncover regulators whose activity may not correlate directly with their expression.

### Integrating spatial and pseudotemporal information improves regulatory inference

Spatial and pseudotemporal information capture complementary aspects of regulatory activity. Spatial information reflects tissue architecture and reveals how a cell’s location shapes the signals it receives, the cell–cell interactions it engages in, and the functional state it adopts^79^. However, spatial variation alone cannot determine whether regional heterogeneity is relevant to the biological processes of interest. Pseudotemporal ordering provides this dynamic context by identifying changes associated with progression along a biological trajectory. By integrating both dimensions, BART-spatial distinguishes trajectory-relevant regulatory signals from unrelated spatial variation and provides a more complete view of how regulatory programs change across tissue space and biological processes.

In the mouse embryonic brain dataset, our analysis highlighted KLF4, CREB1, LEF1, MYCN, E2F3, all of which have established functions in radial glia differentiation during mouse embryonic development. Importantly, these regulators were recovered only when spatial variability was incorporated. They were identified by the full BART-spatial framework but not when inference was restricted to pseudotime related genes alone (Figure S3a). Because TVFs are defined solely from pseudotemporal changes in gene expression, this comparison indicates that trajectory information alone is insufficient to capture the regulatory activity of all active TRs. By integrating spatial and pseudotemporal signals, BART-spatial can therefore identify cell state-specific TR activity that is decoupled from transcript abundance and may be missed by conventional trajectory-based analyses.

Our benchmarking in the OSCC dataset further supports the necessity of integrating both spatial and pseudotemporal information (Figure 2a, b). Although analyses based on either TVFs or SVFs alone recovered part of the 15 known functional TRs, neither approach recovered all of them. For example, SOX2, TP73, TCF4, and GRHL3 were only prioritized when using the overlapped feature set (the intersection of SVFs and TVFs), and missed by either TVF-only or SVF-only approach (Figure S1a-f). Moreover, using the overlapped features, all known functional TRs consistently had scaled ranks higher than 0.75 and were more clearly separated from other TRs (Figure S1g). In contrast, the TVF-only and SVF-only analyses produced less consistent rankings across the known regulators (Figure S1g).

The improvement by using BART-spatial with overlapped features likely increased the signal-to-noise ratio. TVFs alone may include genes whose expression reflect side cellular processes in the trajectory of interest or stochastic variation, which can obscure signals from truly relevant regulators^80,81^. Whereas SVFs alone may highlight genes with strong local variable expression without distinguishing whether that variation is functionally relevant to the biological trajectory of interest. Restricting the analysis to features supported by both spatial and pseudotemporal variation enriches for signals that are simultaneously region-specific and trajectory-relevant, thereby improving the recovery and prioritization of functional TRs.

### Moran’s I enables robust SVF detection in spatial ATAC-seq

A key technical advantage of BART-spatial is its use of Moran’s I statistic to identify SVFs. Moran’s I quantifies spatial autocorrelation in a distribution-free manner, making minimal assumptions about the underlying data^17,82,83^. Moran’s I is computationally efficient, less prone to overfitting, and robust to varying data distributions across tissues or modalities^17,82,83^. This assumption-light approach is particularly useful when applying BART-spatial to spatial epigenomics data, where chromatin accessibility signal is sparser than gene expression and may not follow the same statistical properties that expression-based models are designed to handle^84^. Moran’s I’s flexibility makes BART-spatial one of the few frameworks that can be robustly applied to spatial ATAC-seq data.

We evaluated several existing tools (SpaTrack, SPARK^85^, SPARK-X^86^, SPADE^87^, SpaGene^88^, MERINGUE^89^, singleCellHaystack^90^) designed to detect SVFs from spatial transcriptomics data but most failed for spatial ATAC-seq data. However, Moran’s I-based detection in BART-spatial yields biologically meaningful spatially variable peaks from spatial ATAC-seq data. From the mouse embryo spatial ATAC-seq dataset, BART-spatial identified 581 SVFs out of 32,343 ATAC-seq peaks (p-value < 0.05, deviation of Moran’s I from expectation > 0.1), while other methods detected fewer than 200 peaks under the same p-value threshold (Figure S4a). SpaTrack produced 39 and 30 SVFs with decreased accessibility and increased accessibility along the trajectory, respectively (Figure S4b-d). Notably, SPARK and SPARK-X each took more than five hours running without producing converged results and were therefore excluded from further comparison. Although SPADE initially reported 1,752 peaks, none passed the p-value cutoff of 0.05.

We next evaluated whether the detected SVFs were associated with the developmental trajectory. For each method, we calculated the Pearson correlation between the average peak accessibility and pseudotime at the spot level. We evaluated both the complete set of BART-spatial SVFs and the subset uniquely detected by BART-spatial. Both sets showed the strongest negative correlations with pseudotime among the methods compared (Figure S4e, f). Steiger’s Z-test further showed that the correlation obtained using BART-spatial SVFs was significantly stronger than those obtained using MERINGUE, singleCellHaystack, and SpaGene. (Figure S4e). BART-spatial-unique SVFs also showed significantly stronger correlations than features identified by singleCellHaystack and SpaGene. (Figure S4f).

The spatial accessibility patterns further supported the biological relevance of the detected features. BART-spatial SVFs showed strong accessibility enrichment in radial glia regions, followed by decreasing accessibility along the differentiation trajectory (Figure S4g, h). Similar but weaker patterns were observed for SpaTrack, MERINGUE, and SpaGene, whereas singleCellHaystack did not recover the same pattern. (Figure S4h, j–n). Notably, BART-spatial uniquely identified 496 SVFs that were not detected by any other method. These features also showed marked enrichment of accessible peaks in radial glia regions (Figure S4h, i). Taken together, Moran’s I enables BART-spatial to detect spatially variable chromatin-accessibility features more reliably than existing methods developed primarily for spatial transcriptomics.

### Potential limitations and perspectives

While BART-spatial demonstrates strong capabilities, there are important limitations to consider. First, the BART-spatial framework inevitably relies on user-defined biological inputs, such as the choice of trajectory of interest, region annotations, or cell type labels. Errors or biases introduced during these steps due to subjective labeling, misaligned clusters, or suboptimal trajectory construction can propagate through the pipeline, potentially influencing TR predictions. BART-spatial cannot automatically calibrate or correct user input errors, so careful curation and biological validation remain essential. Second, although BART-spatial’s reference database is extensive, containing over 13,000 publicly available TR binding profiles, gaps still exist for certain TRs. As a result, BART-spatial may underrepresent regulators with limited or no ChIP-seq data, biasing predictions toward well-studied TRs. Third, the accuracy of BART-spatial’s predictions depends on the quality of the input spatial omics data. High-resolution and well-annotated datasets enable more precise detection of SVFs and accurate pseudotime inference. Conversely, data with low resolution, high sparsity, or batch effects may confound spatial signals, leading to false positives or missed prediction. While Moran’s I is robust to some noise, systematic technical artifacts, such as misaligned tissue sections or inconsistently processed replicates, can still introduce biases that BART-spatial alone cannot fully resolve. Ideally, integrating spatial transcriptomics and spatial ATAC-seq data should further improve regulatory inference by combining transcriptional and epigenetic information. However, spatial ATAC-seq remains technically challenging and less widely available. BART-spatial is therefore uniquely valuable for inferring TR activity from spatial transcriptomics data alone when matched epigenomics data are unavailable.

Despite these limitations, BART-spatial represents an important step forward in the computational analysis of spatial omics data, delivering a flexible framework that bridges spatial heterogeneity and biological processes to decode functional transcriptional regulation. Although we have validated selected predictions with matched spatial epigenomics data or existing literature, additional high-quality, high-throughput spatially resolved experiments will be essential for further establishing the biological relevance of predicted TRs and uncovering new regulatory mechanisms. Overall, BART-spatial provides a versatile, accurate, and broadly applicable approach for decoding transcriptional regulation with spatial resolution.

## Methods

### BART-spatial workflow

BART-spatial supports both spatial transcriptomic and epigenomic data. Required inputs include: (1) a gene-by-cell (or peak-by-cell) count matrix, (2) a data frame containing cell metadata, (3) a data frame of feature annotations, (4) a data frame with spatial coordinates for each cell or spot, and (5) a vector specifying cells of interest.

The workflow begins by subsetting the input matrix to retain only the specified cells of interest. BART-spatial then identifies spatially variable features (SVFs) using Moran’s I statistic, a measure of spatial autocorrelation that quantifies whether similar values (i.e., gene expression or chromatin accessibility) cluster spatially. A positive deviation of Moran’s I from expectation indicates non-random spatial organization, reflecting localized gene regulation during cell differentiation.

In parallel, BART-spatial constructs a trajectory starting from a user-defined cell type. Next, Spearman correlation is used to identify features whose expression/accessibility patterns change consistently along pseudotime. These features are defined as temporally variable features (TVFs). Then, we use the intersection of TVFs and SVFs to define the input for the BART algorithm, which predicts functional transcriptional regulators (TRs) that regulate a given set of genes or genomic regions.

For the RNA mode, the BART algorithm^16^ first converts the input gene set into a cis-regulatory profile represented as a ranked list of union DNase I hypersensitive sites (UDHS), leveraging a large collection of public H3K27ac ChIP-seq datasets through the MARGE framework^18^. Then, BART systematically compares this inferred regulatory profile to the over 13,000 TR ChIP-seq binding profiles. Each TR’s binding profile is mapped onto the UDHS regions and represented as a binary vector. BART quantifies the similarity between the input cis-regulatory profile and each TR’s binding profile using area under the ROC curve (AUC) analysis. Statistical significance is assessed using a Wilcoxon rank-sum test, and a final rank is assigned to each TR by integrating the AUC, p-value, and Z-score.

For the ATAC mode, the BART algorithm directly maps the input genomic regions to the UDHS reference. A regulatory profile is constructed by assigning scores to UDHS based on overlap with accessible regions, bypassing the need for MARGE. The same comparison framework is then applied.

### BART-spatial: Rank aggregation in BART-spatial ATAC mode

Instead of using the overlapping features, the ATAC mode takes the union of SVFs and TVFs as input. The ranking of input genomic regions influences the TR prediction results. Ideally, the input should include all regions across the genome, ranked from most to least significant. In order to maximize the sensitivity of BART-spatial and prioritize functionally relevant regions, we designed an additional rank aggregation strategy to compute a composite rank score for each genomic region. This score integrates four components: (1) Moran’s I statistic; (2) the Spearman correlation coefficient between chromatin accessibility and pseudotime; (3) the scaled rank within the set of SVFs; and (4) the scaled rank within the set of TVFs. SVFs and TVFs are ranked independently based on their Moran’s I deviation values and pseudotime correlation coefficients, respectively. The scaled rank for each feature is defined as the inverse of its rank within the corresponding set (i.e., scaled rank = 1/rank). The final rank score for each region is calculated as (Deviation of Moran’s I from expectation + pseudotime correlation) × (scaled SVF rank + scaled TVF rank).

This formulation, conceptually summarized as *(x + y) × (a + b)*, ensures that genomic regions highly ranked in both SVF and TVF sets receive the highest composite scores. Regions that are top-ranked in only one set are moderately favored, while those low-ranked in both sets receive the lowest scores. Genomic regions are then ordered based on their rank scores and input into the BART algorithm for TR prediction.

### BART-spatial: Trajectory analysis

Trajectories for all spatial transcriptomics data were constructed using the “construct_trajectory” function in BART-spatial. This function included “new_cell_data_set”, “preprocess_cds”, “reduce_dimension”, “cluster_cells”, “learn_graph”, and “order_cells” functions of Monocle3^5^ (v1.4.23).

Trajectory inference for the spatial epigenomics datasets was performed using the “addTrajectory” function in ArchR^91^ (v1.0.1). Pseudotime values for individual cells were extracted using the “getTrajectory” function and then integrated into the corresponding Seurat object using the “AddMetaData” function.

Trajectories for all tested datasets, except for the Visium OSCC data, were visualized spatially using the “SpatialFeaturePlot” function of Seurat^92^ (v5.1.0). The inferred pseudotime of Visium OSCC data was visualized with ggplot2.

### BART-spatial: Spatially variable features analysis

By default, spatially variable features were identified using Moran’s I inference in a three-step procedure. First, the “prepare_moran_input” function split the count matrix by feature and generated an object containing individual feature-wise count matrices and spatial coordinates. Second, the “preprocess_data” function scaled and centered both the count matrices and spatial coordinates. Finally, Moran’s I was computed for each gene using the “compute_moran_I” function. Spatial weights, required for the test, were derived based on neighbor relationships. Here, *k* was set to the smaller of 5 or the total number of cells minus one. The “get_moran_result” function computes the deviation of Moran’s I from expectation and output statistically significant features based on user selected cutoff.

Alternatively, BART-spatial provides the option to identify spatially variable features using either SPARK-X or a KNN-based method. The SPARK-X algorithm, specifically developed for spatial transcriptomics data, is detailed in Sun et al., Genome Biology (2021)^86^. SPARK-X is reported to have the best performance by multiple benchmarking studies. However, it is important to note that SPARK-X may not perform well on spatial epigenomics data due to the increased sparsity.

The KNN-based method was implemented in the “run_knn_spatial” function and consisted of two main steps. First, a pairwise distance matrix was computed using the spatial coordinates of all cells, followed by the identification of each cell’s *k* nearest neighbors, excluding the cell itself. The parameter *k* was user-defined and must be smaller than the total number of cells. In the second step, an importance score was calculated for each feature based on the expression matrix. For each feature, expression values were split into two matrices: a self-expression vector representing each cell and a neighbor-expression matrix representing the average expression values of its *k* neighbors. Two methods were supported for computing the gene-level importance score: (1) Correlation method: For each cell, the Spearman correlation coefficient between the gene’s expression in the cell and its neighbors was computed. The final importance score for the feature was defined as the average absolute correlation across all cells. (2) Variance method: For each cell, the variance of the gene’s expression among its neighbors was computed. The importance score for the gene was then defined as the average variance across all cells. Users can select important spatially variable features based on their importance score.

### BART-spatial: Upstream and downstream results integrative analysis

The ranks of TRs in the upstream and downstream prediction lists were converted to their reciprocal values (1/rank), such that higher-ranked TRs received larger scores. TRs identified as significant in either the upstream or downstream list (based on user-defined p-value cutoffs) were assigned a temporary score calculated as the difference between their scaled ranks: (1 / upstream rank) – (1 / downstream rank). TRs that did not meet statistical significance criteria in either list were excluded, ensuring that the final integrative output includes only important TRs. The remaining TRs were re-ranked based on their temporary scores. A positive score indicates upstream regulatory activity, while a negative score reflects downstream activity. A higher absolute score means a higher activity of that TR.

### Spatial omics data processing

#### Visium HPV-negative oral squamous cell carcinoma (OSCC) data processing

This study uses the processed Visium data provided by Aurora and Cao et al., which can be downloaded at https://doi.org/10.6084/m9.figshare.20304456.v1. The spatial distribution of OSCC states, expression of corresponding marker genes, average expression of input genes to BART algorithm was visualized with ggplot2. To improve contrast and reduce the influence of outliers, the upper limit of the color scale was capped at the 99th percentile of expression values.

#### Visium HD mouse small intestine data processing

The Space Ranger-aligned Visium HD data were downloaded from the 10x genomic website at https://www.10xgenomics.com/datasets/visium-hd-cytassist-gene-expression-libraries-of-mouse-intestine. The processed spatial data were analyzed using R software (v.4.4.1) package Seurat (v5.1.0). “008um” resolution was used for this study and spots with low number of features (< 100) were filtered out. After quality control, we followed the standard pipeline of Seurat to process the spatial data, including normalization, scaling and principal component analysis (PCA). To obtain cell type annotation, we projected the labels from a single-cell RNA-seq dataset^36^ to this spatial data using “FindTransferAnchors” and “TransferData” commands. Before projection, we run PCA reduction on the reference scRNA-seq data. For the purpose of this study, we further extract cells with labels containing “Enterocyte” to construct a trajectory for enterocyte differentiation. Cells labelled as “Enterocyte.Progenitor”, “Enterocyte.Progenitor.Late” and “Enterocyte.Progenitor.Early” were grouped as one stage termed “Enterocyte_progenitor”. Similarly, cells labeled as “Enterocyte.Immature.Distal” and “Enterocyte.Mature.Distal” were re-named as “Enterocyte_immature” and “Enterocyte_mature”, respectively. To enable a more focused analysis and better visualization, an area of interest was defined by subsetting the data to include only spots with x-coordinates between 13,500 and 15,000 and y-coordinates between 10,000 and 12,500 using the “subset” function. All spatial maps demonstrating the cell type composition were generated with the “SpatialDimPlot” function in Seurat. Gene expression for key markers was visualized using ggplot2 (v3.5.1). To improve contrast and reduce the influence of outliers, the upper limit of the color scale was capped at the 99th percentile of expression values.

#### Atera human breast cancer data processing

The Space Ranger-aligned Atera data were downloaded from the 10x genomic website at https://www.10xgenomics.com/datasets/atera-wta-ffpe-human-breast-cancer. The processed spatial data were analyzed using R software (v.4.4.1) package Seurat (v5.1.0) following image-based spatial data analysis workflow. Cell type annotations were provided with the Space Ranger-aligned data. Cells labeled as “CAFs, DCIS Associated” or “CAFs, Invasive Associated” within the region x = 0–5000 and y = 7500– 10000 were selected for BART-spatial analysis. All spatial maps demonstrating the cell type composition were generated with the “ImageDimPlot” function in Seurat. Pseudotimes on spatial maps were plotted with “ImageFeaturePlot” function in Seurat.

#### Spatial RNA-ATAC-seq E13 mouse embryo data processing

The processed Seurat object of spatial epigenome-transcriptome co-sequencing E13 mouse embryo data was downloaded from the UCSC Brain Spatial Omics portal at https://brain-spatial-omics.cells.ucsc.edu. Cell type annotations were assigned via label transfer from the Mouse Organogenesis Cell Atlas (MOCA)^93^, following the established pipeline^8^. For the purposes of this study, cells labeled as “radial glia” and “postmitotic premature neurons” were extracted to reconstruct a neuronal differentiation trajectory, as outlined in the same reference. All spatial maps demonstrating the cell type composition and marker gene expression were generated with the “SpatialDimPlot” function in Seurat.

#### Spatial epigenomics E13 mouse embryo data processing

Three spatial epigenomics datasets from E13 mouse embryos were analyzed in this study: (1) the ATAC component of the Spatial RNA-ATAC-seq dataset, (2) replicate 1 and (3) replicate 2 of spatial ATAC-seq. Processed Seurat objects corresponding to these datasets were obtained from the UCSC Brain Spatial Omics portal at https://brain-spatial-omics.cells.ucsc.edu. Raw count matrices were downloaded from GEO (GSE205055) and imported into ArchR (v1.0.1). Data processing, including normalization, dimensionality reduction, and peak calling, was conducted following the established pipeline^8^. Cell type annotations from MOCA were transferred to the processed Seurat objects using the same label transfer method described in the previous section and subsequently added to the corresponding ArchR objects. The purpose of constructing ArchR objects was to enable trajectory inference, as ArchR is specifically designed to handle the sparsity and complexity of single-cell level epigenomics data more effectively than other frameworks.

### Motif analysis

Genome-wide KLF4 motif sites were generated using FIMO^94^ with HOCOMOCO^95^ database and converted as a bed file. Then, to identify peaks containing KLF4 motifs, we overlapped KLF4 motif sites with all peaks measured in the spatial epigenomics data from the Spatial RNA-ATAC-seq dataset. Both sets of genomic regions were first converted into GRanges objects using the “GRanges” function, and overlaps were identified using the “findOverlaps” function from the GenomicRanges package (v1.58.0) with default parameters. For each spatial spot, raw accessibility counts were averaged across all KLF4 motif-containing peaks to derive a KLF4 motif-associated accessibility score. Spots were grouped as radial glia or postmitotic premature neurons according to their cell-type annotations. The spatial distribution of KLF4 motif-associated accessibility was visualized using ggplot2, with the upper limit of the color scale capped at the 90th percentile of accessibility values to enhance contrast and reduce the impact of outliers. Differences in KLF4 motif-containing peak accessibility between radial glia and postmitotic premature neurons were visualized using boxplots. Statistical significance was assessed using a one-sided Wilcoxon rank-sum test to determine whether KLF4 motif-associated accessibility was higher in radial glia. Each point in the boxplot represents one spatial spot.

To assess global motif-associated chromatin accessibility, pseudobulk ATAC-seq profiles were generated separately for radial glia and postmitotic premature neurons by pooling all raw ATAC fragments from spots assigned to each cell type. Peaks were called independently for each pseudobulk population using MACS2 (v 2.2.7.1), and the resulting peak sets were merged to create a union peak set.

For unbiased motif enrichment analysis, fragment counts within each merged peak were normalized by total pseudobulk read depth. Peaks with an absolute log2 fold change greater than 1.6 between radial glia and postmitotic premature neurons were classified as cell-type-enriched accessible regions. These regions were then mapped back to the corresponding cell-type-specific MACS2 narrow peaks, and 600-bp regions centered on the MACS2 summits were used as input for SeqPos^96,97^ motif enrichment analysis.

To determine whether motif enrichment was accompanied by increased local chromatin accessibility at TF binding sites, genome-wide KLF4/CREB1/LEF1/MYCN/E2F3 motif sites that overlapped the union ATAC peaks were retained for downstream analysis. For each motif, Tn5 insertion sites were inferred from fragment ends and counted within ±250 bp of the motif center using 10-bp bins. Motifs on the negative strand were reversed so that all motif-centered profiles were aligned to a common orientation. For each cell type, insertion counts were averaged across motif sites and normalized by the total number of pseudobulk fragments and bin size. The resulting motif-centered accessibility profiles were then compared between radial glia and postmitotic premature neurons for each TF.

### Benchmarking analysis

#### Comparison of TR prediction tools for spatial omics data

The same pipeline was performed on OSCC Visium data, mouse intestine Visium HD data and mouse embryo spatial transcriptomics data.

For SpaTrack (v0.1.1), variable genes were identified using the “ptime_gene_GAM” function. Genes were selected based on the following criteria: “model_fit” > 0.05, “fdr” < 0.05, and “pvalue” < 0.05. To ensure consistency in identifying upstream regulators, genes labeled with a “decrease” expression pattern by SpaTrack were used for upstream-active TR inference, and genes with “increase” pattern were used to predict downstream-active TRs. SpaTrack’s TR prediction was performed using “Trainer”, “get_dataloader”, and “network_df” functions. The reference TR was defined using the list of human TRs provided in the SpaTrack tutorials. As SpaTrack does not include built-in common statistical filters (e.g., p-values) for predicted TRs, all inferred TRs were considered and included in the comparative analysis.

For SCRIPro (v1.1.12), the raw gene expression matrix was processed using the “enrich_rna” function with default parameters. This function groups cells/spots into SuperCells and infers TR activity at the SuperCell level. The output was one TR x SuperCell matrix showing TR activity score and a corresponding p-value matrix. Then, each SuperCell was assigned a state label (upstream or downstream) based on the dominant cell type within each SuperCell and ranked them by TR activity scores. One cell type could be represented more than one SuperCell, so the average of a TR’s activity scores in all SuperCells within a state were taken as its activity score in that state. Next, the prediction results were further separated based on state label and individually ranked based on the averaged TR activity scores at upstream or downstream list.

Two benchmarking metrics were used to evaluate TR prediction performance: (1) Number of experimentally supported functional TRs found in top 25 predictions and (2) Scaled rank of the functional TRs. For the first metric, only the upstream list was used for comparison in OSCC Visium dataset as most known functional TRs appear in the upstream predictions for all methods. In the other two datasets, both upstream and downstream prediction lists were used. Different sets of known functional TRs were compiled based on published experimental evidence supporting their roles in each system of the dataset (Table S1).

To standardize rankings across methods, each TR’s raw rank was rescaled using the formula: scaled rank = (max rank−TR rank) / (max rank - min rank). This transformation yields values between 0 and 1, with higher values indicating stronger predicted regulatory activity. A one-sided Wilcoxon test was used to test whether the scaled ranks of known functional TRs were significantly higher than those of non-functional TRs (alternative hypothesis: functional TRs have higher ranks). Statistical significance was annotated with asterisks (*p* < 1e-5 as *****, *p* < 1e-4 as ****, *p* < 1e-3 as ***, *p* < 1e-2 as **, *p* < 0.05 as *, and *p* > 0.05 as NS), and results were visualized using side-by-side boxplots generated with ggplot2.

### Comparison of SVF detection tools for spatial epigenomics data

Six tools were benchmarked for SVF detection on the ATAC component of the spatial RNA– ATAC-seq co-profiling dataset, including BART-spatial. All methods were provided with the same inputs: a raw peak–by–spot accessibility matrix and spatial coordinates of each spot.

SVFs were identified as follows: MERINGUE (v1.0) using the functions “getSpatialNeighbors,” “getSpatialPatterns,” and “filterSpatialPatterns” with default cutoffs; SpaGene (v0.1.0) using the “SpaGene” function with SVFs filtered at adjusted *p* < 0.05; singleCellHaystack (v1.0.2) using the “haystack” function with SVFs filtered at –log(*p*adj) > –log(0.05); SPADE (v0.99.0) using “SPADE_norm,” “SPADE_estimate,” and “SPADE_test” with SVFs filtered at adjusted *p* < 0.05; and SpaTrack using the pipeline described in the previous section.

Two evaluation metrics were applied: (1) average accessibility of SVFs in radial glia versus postmitotic premature neurons; and (2) correlation with ArchR-inferred pseudotime. For the first metric, raw SVF accessibility was averaged per spot, and spots were assigned to radial glia or postmitotic premature neurons based on cell type labels. Differences between the two groups were visualized using scatter plots qualitatively and boxplots quantitatively, with statistical significance assessed by a two-sided Wilcoxon test. For the second metric, Pearson correlations were computed between pseudotime and the average accessibility of all SVFs at the single-spot level. To assess differences among correlation coefficients, we applied Steiger’s Z-test. This choice was necessary because both paired t-tests and Fisher’s r-to-z tests assume independence between correlations, which is violated here as all correlations share the same pseudotime series. In particular, paired t-tests require independent paired samples, and Fisher’s r-to-z assumes correlations are based on independent observation pairs. Neither holds when a common variable is present. Steiger’s Z-test, however, is specifically designed for comparing two dependent correlations that share one variable in common. It evaluates whether the correlation between variable A and a common variable B is stronger than the correlation between variable C and the same variable B. In our case, the alternative hypothesis was that the correlation between pseudotime and the average accessibility of BART-spatial SVFs is stronger than that between pseudotime and SVFs identified by other methods, while the correlation between the two sets of SVFs themselves was included as a background adjustment.

### Gene Ontology (GO) pathway enrichment analysis

All Gene Ontology (GO) pathway enrichment analyses were conducted using clusterProfiler^98^ (v4.14.4) on both the input gene sets to BART-spatial and the predicted downstream targets of BART-spatial-predicted transcription factors. The downstream target relationships were obtained from the TRRUST database. Gene symbols were first converted to ENTREZIDs using the “select” function. Enrichment analysis was performed with the “enrichGO” function, applying a p-value cutoff of 0.1. GO terms relevant to the trajectory of interest were selected for visualization using ggplot2 (v3.5.1).

### Correlation analysis

BART-spatial output a ranked list of TRs, ranging from 1 to 566 for mouse data and 1 to 918 for human data. Each TR rank was subtracted from the maximum rank and divided by the range (maximum minus minimum), resulting in a scaled rank between 0 and 1, where values closer to 1 indicate higher predicted TR activity. Spearman correlation tests were performed between scaled ranks of TRs derived from different BART-spatial input datasets using the “cor.test” function (method = ’spearman’). Correlation coefficients were visualized using ggplot2 (v3.5.1) as dot plots.

## Code Availability

The source code for BART-spatial and all the scripts used in this study are available at https://github.com/zang-lab/BARTsp

## Acknowledgements

This work is supported by U.S. National Institutes of Health (NIH) grant R35GM133712 (C.Z.).

---

**Supplementary Figure S1.**
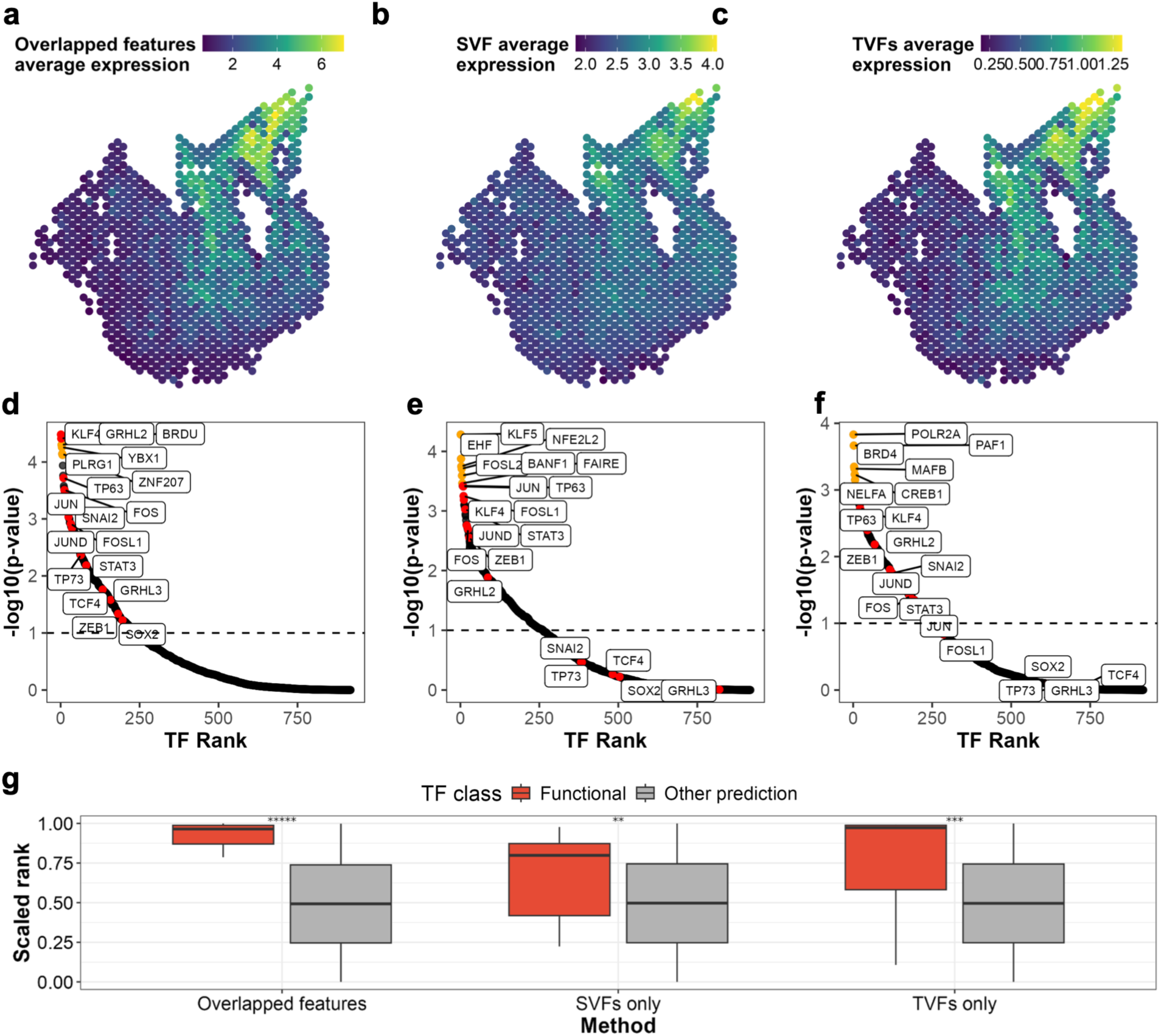
Both pseudotemporal and spatial information are important. (**a**-**c**) Averaged expression of overlapped features between TVFs and SVFs (**a**), all TVFs (**b**), and all SVFs (**c**). (**d**-**f**) BART-spatial prediction results for upstream-active TRs using overlapped features between TVFs and SVFs (**d**), all TVFs (**e**), and all SVFs (**f**). Highlighted in yellow are the top 6 TRs predicted by BART-spatial and highlighted in red are known marker TRs in OSCC development with literature support. (**g**) The scaled ranks of known functional TRs against all the other TRs in the upstream prediction results by BART-spatial using overlapped features between TVFs and SVFs, all TVFs, and all SVFs. *****, p < 10^-5^; ****, p < 10^-4^; NS, non-significant, by one-sided Wilcoxon test.

**Supplementary Figure S2.**
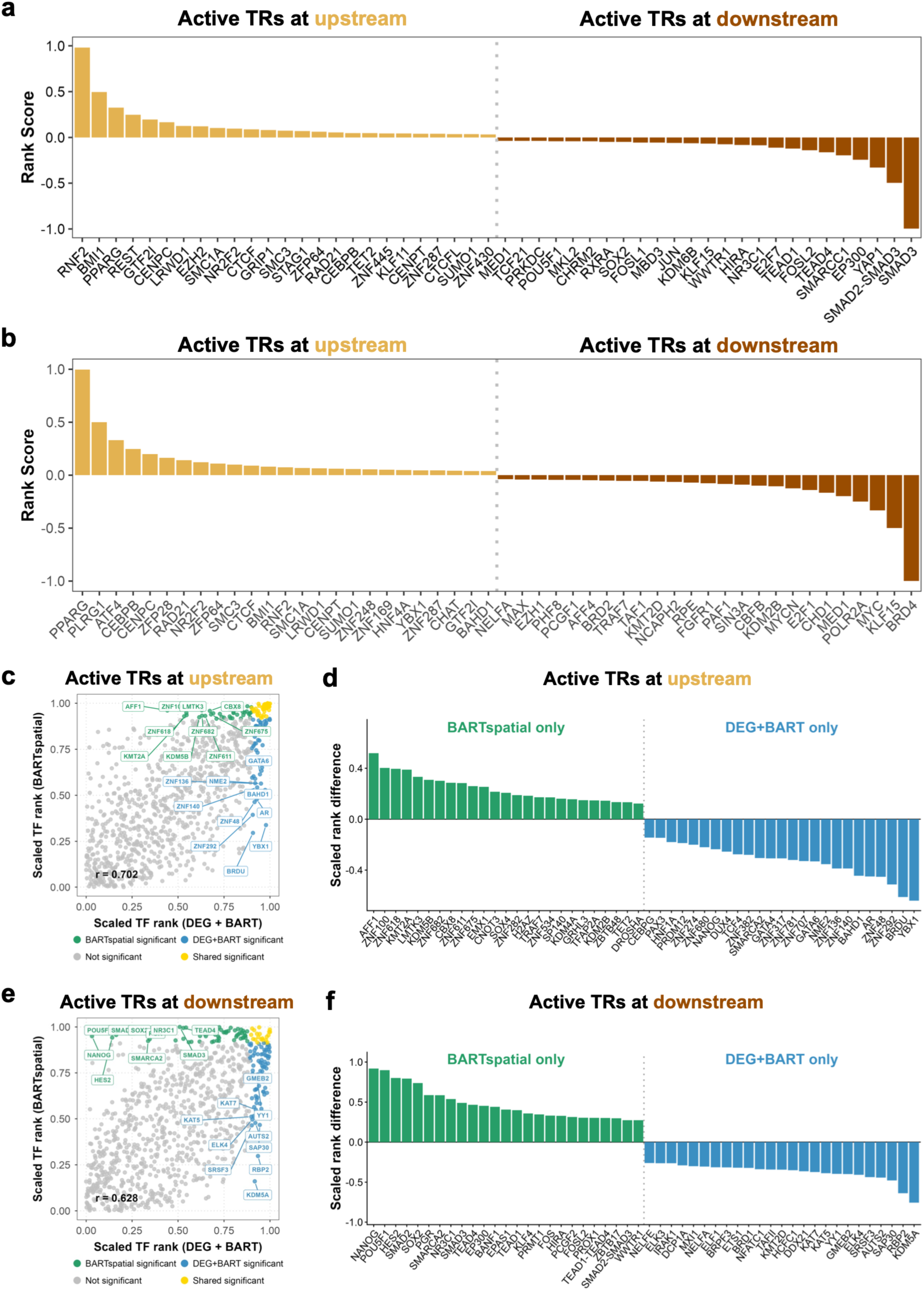
BART-spatial predicts more functionally relevant TRs than DEG+BART. (**a**) Integration of BART-spatial downstream and upstream predictions, showing the top 25 TRs from each of the downstream and upstream TR lists. (**b**) Integration of DEG+BART downstream and upstream predictions, showing the top 25 TRs from each of the downstream and upstream TR lists. (**c**, **d**) Comparison of upstream-active TRs identified by BART-spatial and canonical DEG+BART analysis. TRs significant only by BART-spatial, only by DEG+BART, by both methods, or by neither method are indicated by different colors. (**e**, **f**) Comparison of downstream-active TRs identified by BART-spatial and canonical DEG+BART analysis. TRs significant only by BART-spatial, only by DEG+BART, by both methods, or by neither method are indicated by different colors.

**Supplementary Figure S3.**
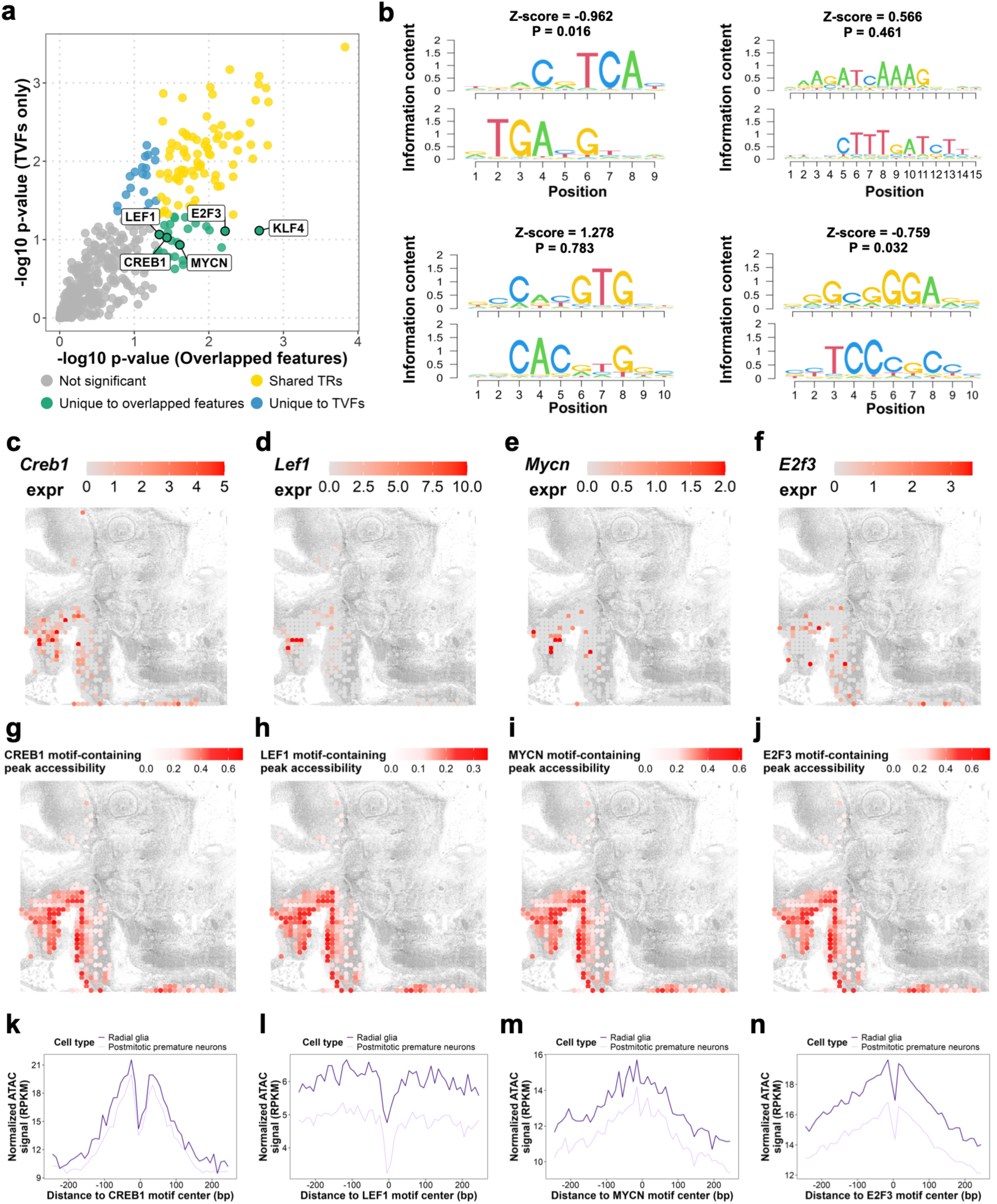
Expression of *Creb1, Lef1, Mycn, and E2f3* and accessibility of their motif-containing peaks demonstrate a similar spatial pattern as KLF4. (**a**) Correlation between TR scaled rank inferred from BART-spatial using pseudotime related genes (TVFs) only and that using overlapped genes. KLF4, CREB1, LEF1, MYCN and E2F3 only appear in the results using overlapped genes and are highlighted in red. (**b**) CREB1 (top left), LEF1 (top right), MYCN (bottom left), E2F3 (bottom right) motif identified from differential ATAC-seq peaks with higher accessibility in radial glia. (**c**-**f**) Spatial map showing expression of *Creb1* (**c**), *Lef1* (**d**), *Mycn* (**e**), *E2f3* (**f**). (**g**-**j**) Chromatin accessibility of motif-containing ATAC-seq peaks for CREB1 (**g**), LEF1 (**h**), MYCN (**i**), E2F3 (**j**), visualized on the spatial map. (**k**-**n**) Aggregate chromatin accessibility profiles centered on CREB1 (**k**), LEF1 (**l**), MYCN (**m**), E2F3 (**n**) motif sites within ATAC-seq peaks.

**Supplementary Figure S4.**
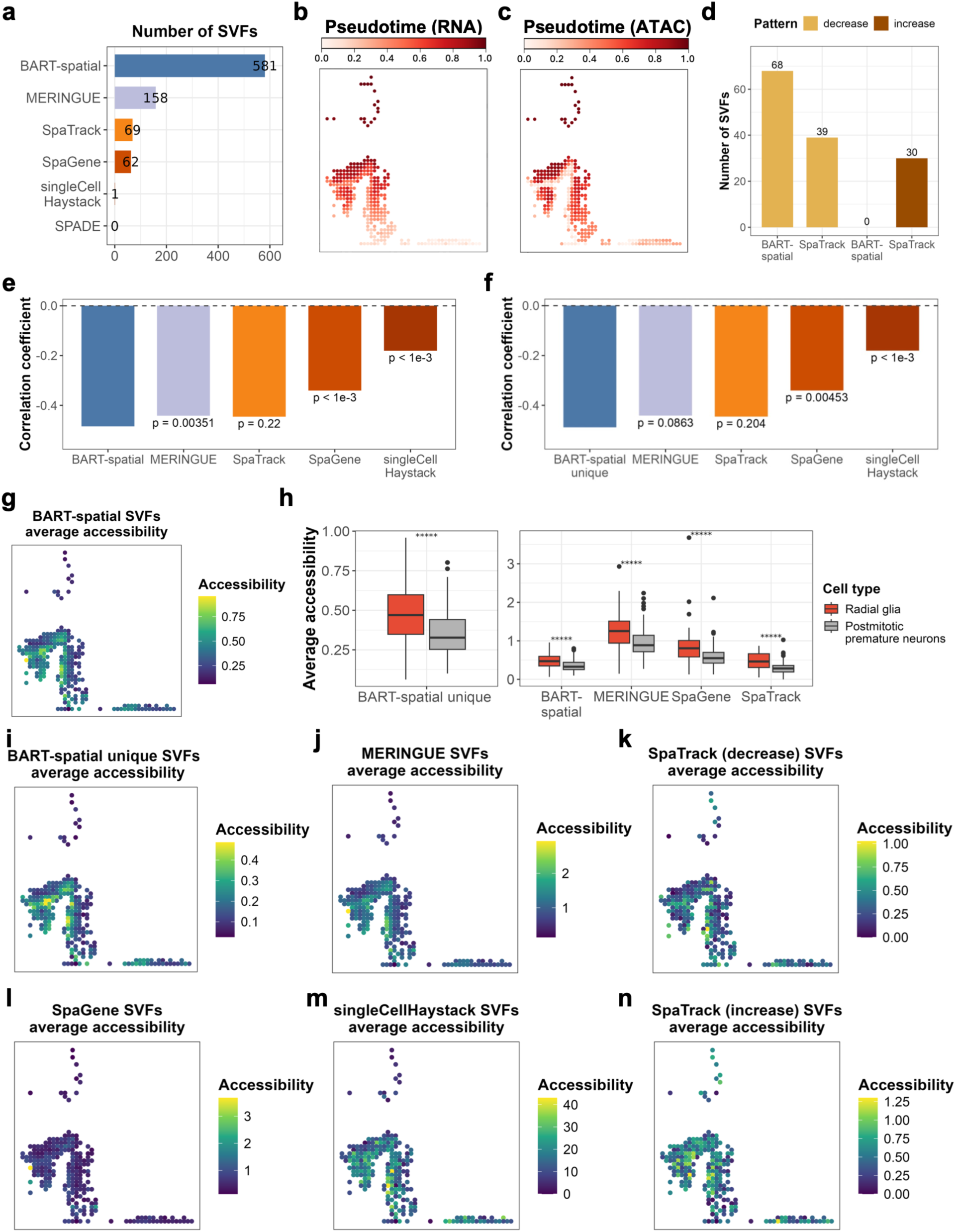
Moran’s I-based BART-spatial finds more biologically meaningful SVFs compared to other methods. (**a**) The number of spatially variable peaks (SVFs) identified by BART-spatial, MERINGUE, SpaGene, singleCellHaystack and SPADE. (**b**, **c**) Pseudotime calculated by SpaTrack on spatial map calculated using RNA part (**b**) and ATAC part (**c**) of the co-profiling data. (**d**) The number of spatially variable peaks identified by BART-spatial and SpaTrack, separated by downstream and upstream. (**e**, **f**) Pearson correlation between pseudotime and the average accessibility of all the SVFs (**e**) and unique SVFs (**f**) identified by BART-spatial, MERINGUE, singleCellHaystack, SpaGene, and SpaTrack using genes with decreasing pattern, as well as unique SVFs identified by BART-spatial. P-values are calculated based on Steiger’s Z-test with average accessibility of BART-spatial derived SVFs as baseline. (**g**) Chromatin accessibility of BART-spatial-identified SVFs visualized on the spatial map. (**h**) Difference in average accessibility of unique SVFs (left) and all SVFs (right) identified by BART-spatial in radial glia and postmitotic premature neurons. *****, p < 10^-5^; NS, non-significant by two-sided Wilcoxon test. (**i**-**n**) Chromatin accessibility of all SVFs detected by BART-spatial-unique (**i**), MERINGUE (**j**), SpaTrack (decrease) (**k**), SpaGene (**l**), singleCellHaystack (**m**), and SpaTrack (increase) (**n**), visualized on the spatial map.

**Table S1.** Functional TRs with literature support for each benchmarking dataset.

| Dataset | Known functional TRs in this system |
| --- | --- |
| HPV- oral squamous cell carcinoma Visium | GRHL3, TCF4, TP63, TP53, GRHL2, SOX2, KLF4, TP73, SNAI2, ZEB1, FOSL1, STAT3, FOS, JUN, JUND |
| Mouse small intestine Visium HD | HNF4G, HNF4A, HNF1B, GATA6, GATA4, MAF, MAFB, CDX2, HES1 |
| E13 mouse embryo spatial RNA-ATAC-seq co-profiling | PAX6, SOX9, KLF4, FEZF2 (Radial glia/Upstream)<br>NEUROD2 (Postmitotic premature neurons/Downstream) |

## Notes

https://github.com/zang-lab/BARTsp

### Competing Interest Statement

The authors have declared no competing interest.

### Summary of Updates

Figures, Supplementary Figures, Results, and Discussion have been revised to improve clarity and readability.

